# Alpha/Beta oscillations reflect the dynamics of the oculomotor system: a new perspective on subsequent memory effects

**DOI:** 10.1101/2025.07.29.667451

**Authors:** Xiongbo Wu, Tzvetan Popov, Tara Beilner, Nicholas Fearns, Christian Vollmar, Elisabeth Kaufmann, Jan Remi, Tobias Staudigl

## Abstract

Neural activity and eye movements are two well-established predictors of memory performance in humans. Successful memory formation is typically associated with reduced alpha/beta power (i.e., the alpha/beta subsequent memory effect) and an increased number of eye movements. However, the functional relevance of these two memory correlates has primarily been investigated in isolation, leaving their coupling and combined contribution to memory formation largely unknown. Here, we address this gap through four experiments involving simultaneous eye-tracking and scalp or intracranial electroencephalography recordings while participants viewed scenes either freely or under varying levels of visual constraint. Across experiments and cohorts, we identified the degree of visual exploration, rather than subsequent memory performance, as the consistent and robust modulator of alpha/beta power reductions. Going beyond correlations, manipulating eye movements dissociated alpha/beta power reductions from memory. Combined with the finding that saccade parameters predicted the dynamics of alpha/beta activity, our results directly link alpha/beta activity to the oculomotor system. Moreover, we show that the accumulation of saccades over time accounted for the reduced alpha/beta power, resembling the pattern typically observed after stimulus onset. Our work demonstrates that neglecting eye movements results in an incomplete understanding of the alpha/beta subsequent memory effect. Together, the evidence points at alpha/beta activity directly reflecting eye movements, and only indirectly, memory formation. The current study thus bridges two lines of previous research and advocates for new perspectives on the specific interpretation of the alpha/beta subsequent memory effect and on the function of alpha/beta activity in general.

## 1. Introduction

In the quest to understand what best predicts successful memory formation in humans, several strong candidates have been identified. Among those, neural oscillations have been suggested to be a key feature when it comes to episodic memory formation^1–3^. Another robust predictor is visual exploration, which highlights memory formation as an active process constantly shaped by, and interacting with, viewing behaviour^4^. While each predictor has been extensively studied in isolation, the relationship between them and their potential coupling in support of memory formation has been largely overlooked.

Previous studies investigating neural oscillations underlying memory formation consistently report that power modulations in the alpha/beta range (approximately 10–20 Hz) predict subsequent memory performance^5–9^ (for other neural predictors of memory, see, e.g.,^2,3^). Specifically, materials later remembered in a memory test are typically associated with stronger alpha/beta power reductions during the study phase, as compared to materials that were later forgotten, a phenomenon termed as alpha/beta subsequent memory effect (SME)^10^. The functional relevance of this alpha/beta SME has been commonly interpreted as reflecting memory processes, emphasizing that low-frequency desynchronization might be optimal for information coding (see^10–14)^. On the other hand, research that investigates eye movements consistently report that more extensive visual exploration during study promotes memory formation^15^. Specifically, better memory performance at test is shown to be associated with a higher number of eye movements during study^4,16–19^. Moreover, studies indicate that patterns of eye movements made during study re-occur at test^20–22^, suggesting visual exploration forms part of the memory formation process.

Recent studies indicate a close link between eye movements and modulations in alpha/beta neural activity. For instance, alpha power reductions have been shown to be modulated moment-to-moment by eye movements^23,24^ even in the absence of visual input^25^. Furthermore, eye movements transiently lateralize alpha power in posterior brain regions, during tasks that prompt large saccades^26^ as well as small saccades under fixations^27^. Initial evidence supporting such a link during memory formation was provided in a recent study by Popov & Staudigl (2023)^28^, which offered the first, albeit correlational, evidence for a covariation of eye movements and the alpha/beta SME.

In the current study, we go beyond this correlational evidence and hypothesize a driving role of eye movements in shaping the well-documented alpha/beta SME. We conducted four memory experiments with simultaneous recordings of eye movements and scalp or intracranial electroencephalography (EEG). Importantly, we manipulated participants’ eye movements by allowing them to view scenes either freely or under varying degrees of visual constraint, and assessed how these manipulations drove corresponding changes in alpha/beta activity. Across four experiments and cohorts, we consistently found that the degree of visual exploration predicted the magnitude of alpha/beta power reductions, even under attempted fixation where the alpha/beta SME was absent. Furthermore, saccade parameters predicted the dynamics of alpha/beta amplitude, and the accumulation of saccades over time accounted for the overall reduction in alpha/beta power. Together, these findings offer a new perspective on alpha/beta power modulations observed in memory paradigms, and support the view that the dominant rhythm in the human brain reflects oculomotor activity.

## 2. Results

### 2.1. Visual exploration and alpha/beta power covary during free viewing and jointly predict subsequent memory (Experiment 1 - Healthy participants)

Twenty participants (8 females; age range: 18–32 years; M = 23.77) were instructed to freely view 216 scene images (size: 28 x 21 degrees of visual angle (dva); duration: 4 seconds (sec) each) and indicate whether a scene was either an indoor or outdoor scene. After a break, participants’ memory for the scenes was tested (see Methods & Fig. 1A). Scalp EEG and gaze positions were simultaneously recorded during the experiment. On average, participants’ accuracy for indoor/outdoor judgement was at ceiling (M = 97.57%, SD = 2.87%). Participants remembered 70.12% of the scenes (SD = 11.33%). All participants showed a *d’* (d-prime) above chance (M = 1.55, SD = 0.56; Fig. 1A right).

**Figure 1.**
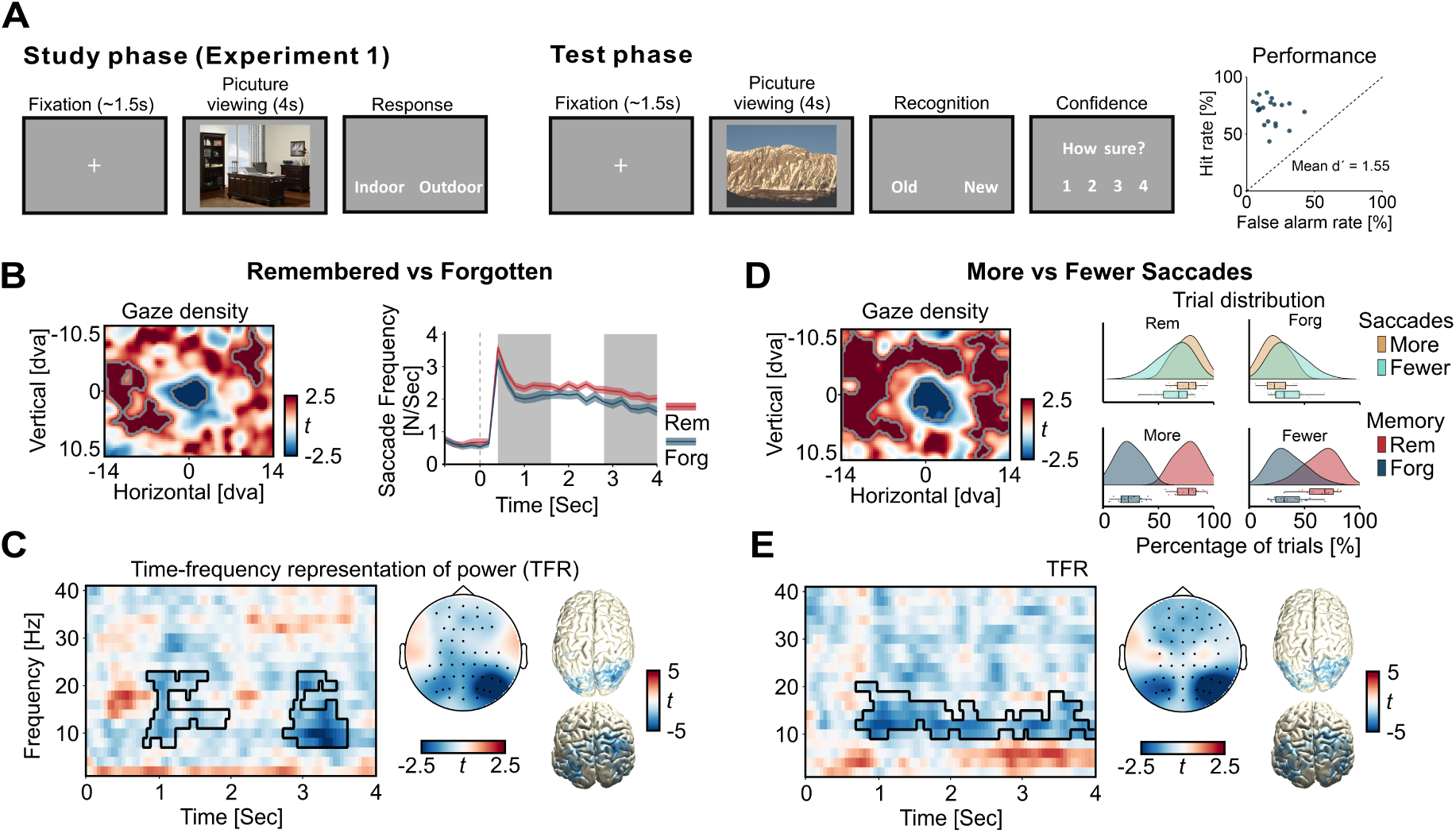
Visual exploration and alpha/beta power covary during free viewing and jointly predict subsequent memory. (**A**) Experimental paradigm for Experiment 1 and memory performance. Each dot represents one participant. (**B**) Increased visual exploration and more eye movements were observed for scenes that were later remembered compared to those that were forgotten. Gaze density contrasts (left) revealed a significant bias towards surrounding scene locations (colour coded in red) and reduced focus on the centre (colour coded in blue) for later remembered as compared to later forgotten scenes. Saccade frequency (right) was higher for later remembered than later forgotten scenes across multiple time windows during viewing. Significant clusters, corrected using a cluster-based permutation test (*p* < 0.05, two-tailed), are indicated by grey contours (left) and shaded grey regions (right). (**C**) Time-frequency representation of power (TFR; averaged across significant sensors marked in the topography) and topography (averaged t-values within the identified significant cluster marked on the TFR) showed reduced alpha/beta power for later remembered compared to later forgotten scenes (*p* < 0.05, two-tailed). Source reconstruction (right) localized this effect to posterior brain regions. (**D**) Visual exploration covaried with alpha/beta power modulation and memory performance. Trials were median-split based on the numbers of saccades during scene viewing. Gaze density contrasts (left) revealed a significant bias towards surrounding scene locations (colour coded in red) and reduced focus on the centre (colour coded in blue) for scenes explored with more as compared to fewer saccades (*p* < 0.05, two-tailed). Rain cloud plots (right) indicate a higher percentage of later remembered trials for scenes viewed with more saccades. (**E**) Scenes explored with more saccades were accompanied by a greater reduction in alpha/beta power centred over the posterior regions of the brain compared to less visually explored scenes (*p* < 0.05, two-tailed).

#### More visual exploration when viewing later remembered vs forgotten images

During scene viewing, participants’ horizontal and vertical gaze positions were tracked across the 4-sec scene presentation and binned into a two-dimensional (2-D) array aligned with the size of the image. After normalizing and smoothing using a 2-D Gaussian filter, the 2-D array for gaze density of each trial was separated by memory performance (i.e., subsequently remembered or forgotten at test phase). A non-parametric cluster-based permutation test was applied to test the difference in gaze density. The analysis revealed a significant bias towards surrounding scene locations and less focus on the centre when viewing scenes that were later remembered as compared to those that were forgotten (Fig. 1B, left; all clusters with *p* < 0.05, two-tailed, corrected for multiple comparisons across space).

Saccade analyses yielded converging results. Participants made significantly more saccades across multiple time windows when viewing later remembered as compared to forgotten scenes (Fig. 1B, right; cluster-based permutation test, all clusters with *p* < 0.05, two-tailed, corrected for multiple comparisons across time). On average, participants made 2.29 saccades per sec (SD = 0.42) viewing later remembered scenes which was significantly more than when viewing later forgotten scenes (M = 1.98, SD = 0.48; paired-sample *t*_(19)_ = 6.25, *p* < 0.001) (for main sequences, saccade durations and direction distributions, see Supplementary Fig. 1).

To further examine the influence of stimulus properties on eye movements and memory, we quantified the visual memorability of each image using MemNet^29^, a pre-trained convolutional neural network (CNN; Supplementary Methods). We then assessed the relationships between image memorability, eye movements, and memory performance. Specifically, we used a generalized linear mixed-effects model (GLMM) to test whether memory performance (binary: remembered vs. forgotten) was predicted by image memorability scores and saccade counts, and a linear mixed-effects model (LMM) to examine whether image memorability scores predicted saccade counts (see Supplementary Methods for details). The analyses revealed that higher image memorability scores significantly predicted better memory performance (*p* = 0.004; Supplementary Fig. 2A). Moreover, when saccade count was included as a fixed effect, a higher number of saccades significantly predicted better memory performance (*p* < 0.001) and improved the overall model fit (*p* < 0.001). In contrast, image memorability did not significantly predict saccade counts during viewing (*p* = 0.58; Supplementary Fig. 2B). Together, these results indicate that although intrinsic image properties influence memory, eye movements play an active contribution to memory formation that is not merely driven by the stimuli themselves.

#### Visual exploration and alpha/beta power jointly predict subsequent memory performance

We next tested whether the neural activity elicited during the study phase differed between later remembered and forgotten scenes and replicated the well-documented alpha/beta subsequent memory effect (SME)^10^. Time-frequency representations (TFRs) of EEG signals were computed time-locked to scene onset, spanning 2–40 Hz in 2 Hz steps. Power estimates were averaged across trials by memory condition and baseline-corrected to the pre-stimulus interval [-1, -0.5] sec.

Consistent with prior findings, we found that later remembered scenes elicited significantly greater power reductions in the alpha/beta range as compared to later forgotten scenes (Fig. 1C, left; *p* < 0.05, two-tailed, corrected for multiple comparisons across time, frequency and sensors). Source reconstruction indicated posterior brain regions underlying the effect (Fig. 1C, right), in line with the spatial distribution commonly reported in the literature^5,10,28,30^.

Next, we explored whether visual exploration covaried with alpha/beta power modulation and memory performance. To do so, we median-split the trials based on total saccade count throughout the 4-sec scene viewing. Gaze density contrasts revealed a strong bias towards surrounding scene locations and less focus on the centre when more saccades were made during scene viewing (Fig. 1D, left; all clusters with *p* < 0.05, two-tailed, corrected for multiple comparisons across space), indicating more visual exploration via more frequent eye movements. Importantly, we found that later remembered trials were for the most part also trials with more saccades, while later forgotten trials where trials with fewer saccades (Fig. 1D, right top). In line with this, trials with more saccades had a stronger tendency to be subsequently remembered (Fig. 1D, right bottom), suggesting that the degree of visual exploration predicted later memory performance.

Notably, we observed a strong alpha/beta power reduction when comparing trials with more to fewer saccades (Fig. 1E, left; *p* < 0.05, two-tailed, corrected for multiple comparisons across time, frequency and sensors), with a temporal and topographical distribution comparable to the observed alpha/beta SME. Scenes explored with more saccades at the study phase elicited significantly stronger alpha/beta power reductions. Source reconstruction localised this effect to the posterior regions of the brain (Fig. 1E right;), very similar to the source reconstruction of the alpha/beta SME.

To investigate whether the observed difference reflected oscillatory desynchronization rather than changes in the aperiodic background activity, we applied the Fitting Oscillations and One-Over-F (FOOOF) algorithm to separate periodic oscillations from aperiodic component^31^ (see Supplementary Methods for details). We then repeated the same statistical comparisons between later remembered and forgotten scenes, as well as between trials viewed with more versus fewer saccades, using the FOOOF-extracted periodic power. The results mirrored those obtained from the non-FOOOFed EEG power analyses, confirming that the observed alpha/beta power reductions reflect oscillatory changes rather than aperiodic signal shifts, in particular in the alpha band (Supplementary Fig. 3A & B). As current study aims to align with the broader alpha/beta literature, which typically reports results based on non-FOOOFed power, we focused on the non-FOOOFed EEG signals for the subsequent analyses to ensure comparability with prior work.

Together, these results from the healthy cohort in Experiment 1 support a functional coupling between visual exploration and alpha/beta power modulation at posterior regions of the brain in service of episodic memory formation. The results replicate and extend recent findings by Popov & Staudigl (2023)^28^, supporting that the relationship between visual exploration and alpha/beta power modulation generalises from MEG to EEG.

### 2.2. Intracranial evidence support greater visual exploration leads to better memory and stronger alpha/beta power reductions (Experiment 2 - Clinical cohort)

Experiment 2 included four drug-resistant epilepsy patients (1 female; age range: 21–36 years; M = 27.25) with intracranial recordings, selected because they had intracranial electrodes implanted in posterior brain regions (Fig. 2B; note that posterior implantations sites are rare in the cohort we recruited from; for detailed electrode localisations see Supplementary Table. 1). The task structure and instructions were identical to those of healthy participants in Experiment 1, with adjustment to trial numbers and block structure to optimize the experiment for clinic settings (see Methods for details). Intracranial EEG and gaze positions were simultaneously recorded throughout the experiment. On average, all four patients performed at ceiling on the indoor/outdoor judgment task (M = 96.18%, SD = 6.74%). Overall, patients remembered 50.69% of the scenes (SD = 26.75%). All patients showed a *d’* above chance (M = 1.19, SD = 0.50).

**Figure 2.**
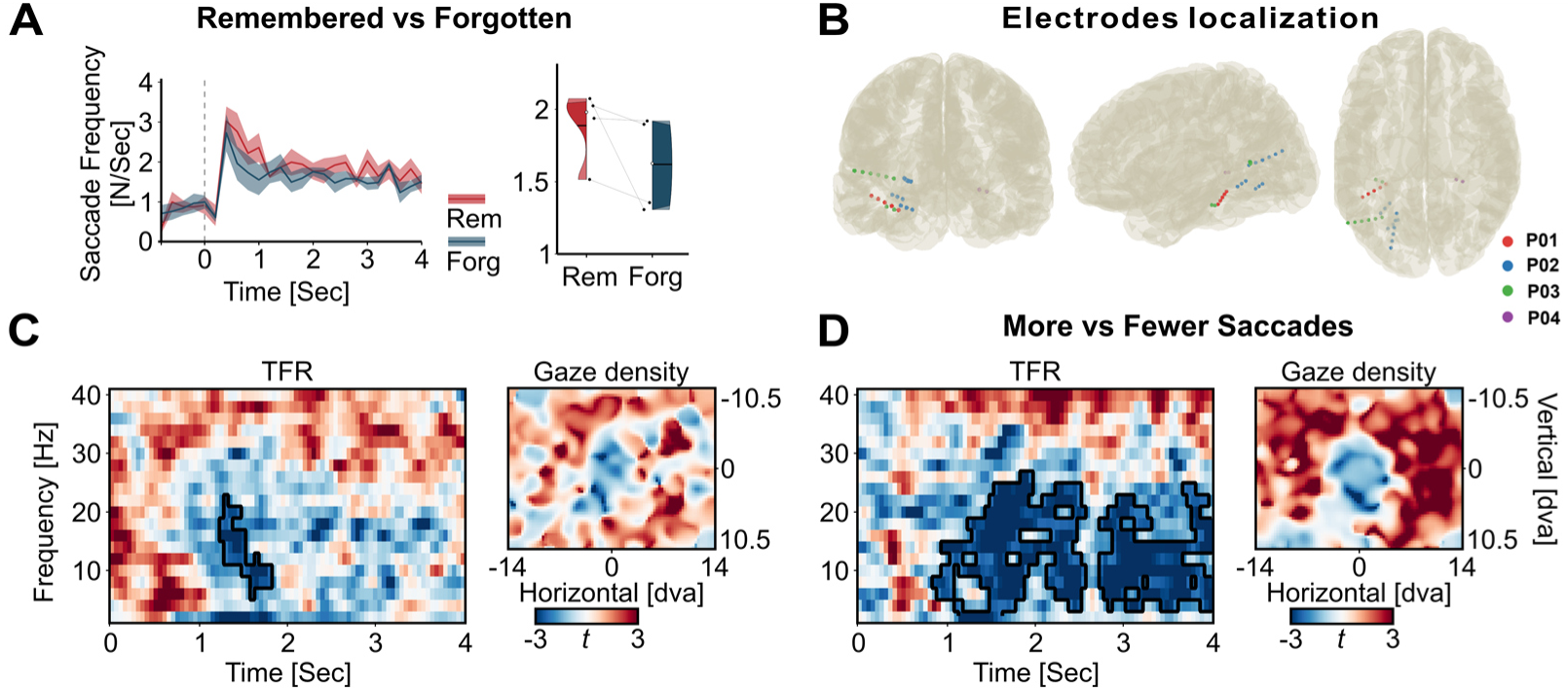
Intracranial data confirms that greater visual exploration leads to better memory and stronger posterior alpha/beta power reductions. (**A**) Saccade frequency across scene viewing (left) in Experiment 2. Each patient (N=4) showed higher average saccade frequency when viewing later remembered compared to forgotten scenes (right). (**B**) Localization of posterior electrodes included in the analyses, shown from back, left and top views. A total of 27 posterior electrodes were recorded across the four patients, resulting in N = 20 bipolar-referenced channels. (**C**) TFR contrasts revealed a reduction in alpha/beta power at posterior electrodes during scene viewing for later remembered as compared to forgotten images (left). Gaze density showed a bias toward surrounding scene locations for later remembered scenes. (**D**) TFR contrasts revealed a prominent reduction in alpha/beta power at posterior electrodes for scenes viewed with more versus fewer saccades (left; trials median-split by saccade count). Corresponding gaze density showed a bias towards surrounding scene locations for later remembered scenes (right). (**C** & **D**) Bipolar-referenced channels (N = 20) were treated as the unit of analysis for TFR comparisons. Significant clusters, corrected using a cluster-based permutation test (*p* < 0.05, two tailed), are indicated by black contours.

#### Intracranial data confirm the findings on scalp EEG

First, we found the same predictive eye movements patterns for subsequent memory in the clinical cohort as observed in the healthy participants. Later remembered scenes were explored with more frequent saccades at the study phase (Fig. 2A, left). All patients showed higher average saccade frequency for later remembered (M = 1.89; SD = 0.25) as compared to forgotten scenes (M = 1.62; SD = 0.33; Fig. 2A, right) (for main sequences, saccade durations and direction distributions, see Supplementary Fig. 1). This pattern was consistent with the gaze density contrast (Fig. 2C, right), where later remembered scenes showed a bias toward surrounding scene locations during viewing as compared to later forgotten ones.

Next, we examined whether posterior intracranial EEG recordings would reveal the alpha/beta SME, and more importantly, whether greater visual exploration would again be associated with stronger alpha/beta power reductions, as observed over posterior scalp regions in the healthy cohort. To do so, TFRs were computed for each bipolar-referenced intracranial EEG channel after preprocessing using the same procedures as with the scalp EEG data (see Methods for details). We treated each channel (N = 20) as the unit of analysis for statistical comparisons. We separated the trials based on subsequent memory (remembered versus forgotten) as well as the degree of visual exploration (more versus fewer saccades via median split).

The results confirmed the alpha/beta SME. Posterior intracranial electrodes showed greater alpha/beta power reductions viewing later remembered as compared to forgotten scenes (Fig. 2C, left; *p* < 0.05, two-tailed, corrected for multiple comparisons across time and frequency). Importantly, a pronounced difference in alpha/beta power reductions was found for scenes explored with more as compared to fewer saccades (Fig. 2D, left; all clusters with *p* < 0.05, two-tailed, corrected for multiple comparisons across time and frequency). This finding conceptually replicates results from the scalp EEG and extends them to direct intracranial recordings, further supporting that greater visual exploration (also illustrated by gaze density contrast displayed on Fig. 2D, right) leads to stronger posterior alpha/beta power reductions.

To summarize, Experiments 1 and 2 showed consistent results in both healthy and clinical cohorts: visual exploration and posterior alpha/beta power reductions co-varied during free viewing of scenes and jointly predicted subsequent memory.

### 2.3. Attempted fixation eliminates alpha/beta subsequent memory effect, while visual exploration modulates alpha/beta power (Experiment 3 - Healthy participants)

In Experiment 3, twenty participants (13 females; age range: 19–33 years; M = 25.05) were instructed to strictly maintain fixation on a centrally displayed red fixation cross (size: 1 x 1 dva) while viewing 216 scenes, each followed by an indoor/outdoor judgment (Fig. 3A, left; see Methods for details). Memory for the scenes was tested in a later phase. Scalp EEG and gaze positions were simultaneously recorded during the experiment. On average, participants’ accuracy for indoor/outdoor judgement was at ceiling (M = 97.36%, SD = 2.30%). Under attempted fixation, participants remembered 64.31% of the scenes (SD = 11.23%), and *d’* indicate all participant performed above chance (M = 1.06, SD = 0.42; Fig. 3A, right). Markedly, memory performance under attempted fixation was significantly lower than free viewing in Experiment 1 (two-sample *t*_(38)_ = -3.12, *p* = 0.003), in line with previous studies^4,32,33^.

**Figure 3.**
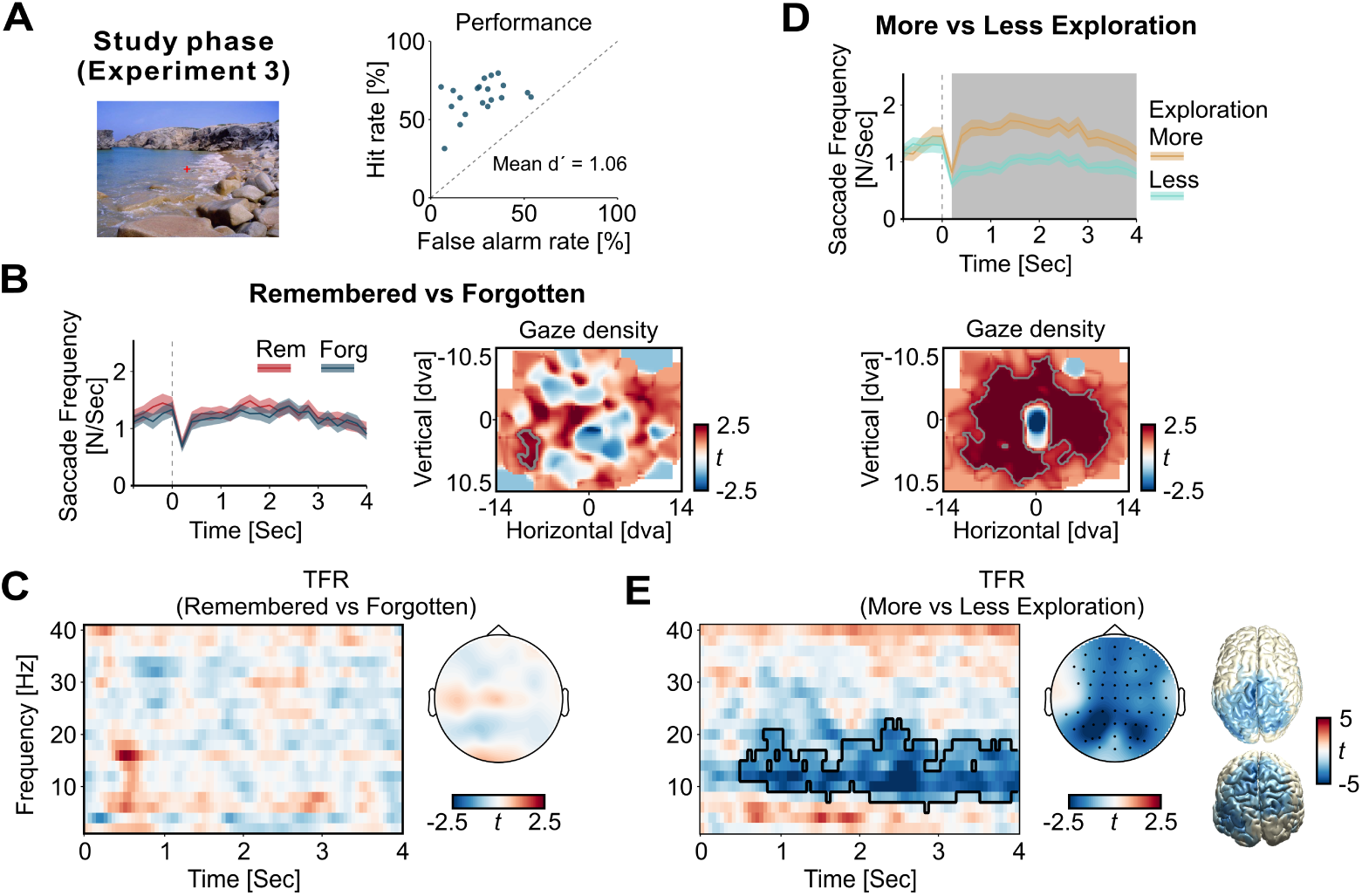
Attempted fixation eliminates alpha/beta subsequent memory effect, while visual exploration modulates alpha/beta power. (**A**) Experimental paradigm for Experiment 3 (left) and memory performance (right). At the study phase, participants were instructed to maintain fixation on a red cross (size: 1 × 1 dva, shown at actual scale relative to the scene) displayed at the centre of the scene throughout the 4-sec viewing period. Trial structure at the study phase and the subsequent memory test were identical to Experiment 1 & 2. (**B**) No significant difference in saccade frequency was found between later remembered versus forgotten scenes during viewing (left). Gaze density contrasts (right) revealed a localized peripheral bias associated with later remembered scenes. (**C**) No significant subsequent memory effect was found in the TFR during the viewing of later remembered as compared to forgotten scenes (Topography showed average t-values within the 10–20 Hz range and the 0.5–4 sec time window). (**D**) Trials median-split by the exploration index (see Methods) showed that scenes with higher visual exploration were associated with significantly more saccades during viewing (top) and a gaze bias towards surrounding scene locations (bottom). (**E**) More visually explored scenes showed a greater reduction in alpha/beta power compared to less visually explored scenes over posterior brain regions. (**B, C, E**) Significant clusters are indicated by grey regions and grey or black contours (cluster-based permutation test, *p* < 0.05, two-tailed).

First, we observed that attempted fixation largely reduced degree of visual exploration throughout scene viewing. On average, participants made 1.21 saccades per sec (SD = 0.43), a frequency comparable to saccade rates commonly reported in human fixation tasks^34–38^. Across subjects, the average saccade size was 1.00 dva (SD = 0.51), with 79.33% (SD = 14.85%) of saccades being smaller than 1 dva—commonly considered as the threshold for microsaccades^37,39^. These results indicate that attempted fixation did not abolish eye movements entirely; instead, participants continued to make fixational eye movements such as microsaccades across stimulus presentation^39^ (for main sequences, saccade durations and direction distributions, see Supplementary Fig. 1).

#### Attempted fixation eliminates alpha/beta subsequent memory effect

When separating the trials based on subsequent memory, we found no significant difference in saccade frequency across time between later remembered and forgotten scenes (Fig. 3B, left). On average, participants made 1.23 saccades/sec (SD = 0.45) when viewing later remembered scenes, not significantly different from the frequency for forgotten scenes (M = 1.17, SD = 0.42; paired-sample *t*_(19)_ = 1.55, *p* = 0.14). Gaze density contrasts (Fig. 3B, right) revealed a localized peripheral bias associated with later remembered images (*p* < 0.05, two-tailed, corrected for multiple comparisons across space). These results indicate an overall comparable degree of visual exploration between the two memory conditions.

We then applied the same analysis procedure as in Experiment 1 to examine the neural activity at the study phase. Importantly, no significant difference was identified when comparing the TFRs between later remembered and forgotten scenes (Fig. 3C), indicating that under attempted fixation, the alpha/beta SME was no longer present.

#### Degree of visual exploration predicts alpha/beta power reduction

Having shown that the alpha/beta power was not associated with subsequent memory under attempted fixation, we next examined whether its modulation by visual exploration would nevertheless persist. Given the overall low saccade frequency under attempted fixation, median-splitting trials by saccade count as in Experiment 1 was not feasible. Instead, we quantified the degree of visual exploration using an exploration index, calculated for each trial as the standard deviation of the normalized and smoothed 2-D gaze density array during scene viewing (see Methods for details and Supplementary Fig. 4 for illustration). Based on this metric, both saccade frequency and gaze density contrast showed consistent patterns (Fig. 3D): more visually explored scenes were associated with significantly higher saccade frequency across viewing (Fig. 3D, top; *p* < 0.05, two-tailed, corrected for multiple comparisons across time) and stronger gaze bias towards surrounding scene locations and reduced central fixation as compared to less visually explored scenes (Fig. 3D, bottom; *p* < 0.05, two-tailed, corrected for multiple comparisons across space).

Importantly, and in contrast to the absent alpha/beta SME in this experiment, we observed significantly reduced alpha/beta power for more visually explored scenes compared to less explored scenes (Fig. 3E left; *p* < 0.05, two-tailed, corrected for multiple comparisons across time, frequency and sensors). This effect was source-localized to posterior brain regions (Fig. 3E right), similar to the spatial distribution observed under free viewing in Experiment 1.

Together, the results from Experiment 3 indicate that the alpha/beta SME no longer persist under attempted fixation when there was minimal difference in visual exploration between later remembered and forgotten scenes. However, when separating the trials based on the degree of visual exploration, stronger reduction in alpha/beta power re-emerged for scenes which were explored more. This suggests a strong link between the alpha/beta modulation and degree of visual exploration, and a less strong association between alpha/beta activity and subsequent memory performance.

### 2.4. Restricting visual exploration modulates the degree of alpha/beta power reductions (Experiment 4 - Healthy participants)

To further investigate the link between visual exploration, alpha/beta power and memory, we conducted Experiment 4, in which we introduced varying degrees of constraints on scene viewing. Thirty-four participants (19 females; age range: 21–35 years; M = 25.97) completed the experiment. Participants were instructed to maintain gaze within a centrally overlaid red circle while viewing 324 scenes, each followed by an indoor/outdoor judgment (Fig. 4A, top). Memory for the scenes was tested in a later phase. The experiment contained three levels of viewing restriction, defined by the diameter of the centrally overlaid red circle: 1, 5, or 21 dva. Trial structure at study phase and test phase was identical to Experiment 1, 2 & 3 (see Methods for details). Scalp EEG and gaze positions were simultaneously recorded during the experiment. On average, participants’ accuracy for indoor/outdoor judgement was at ceiling (M = 97.71%, SD = 1.84%).

**Figure 4.**
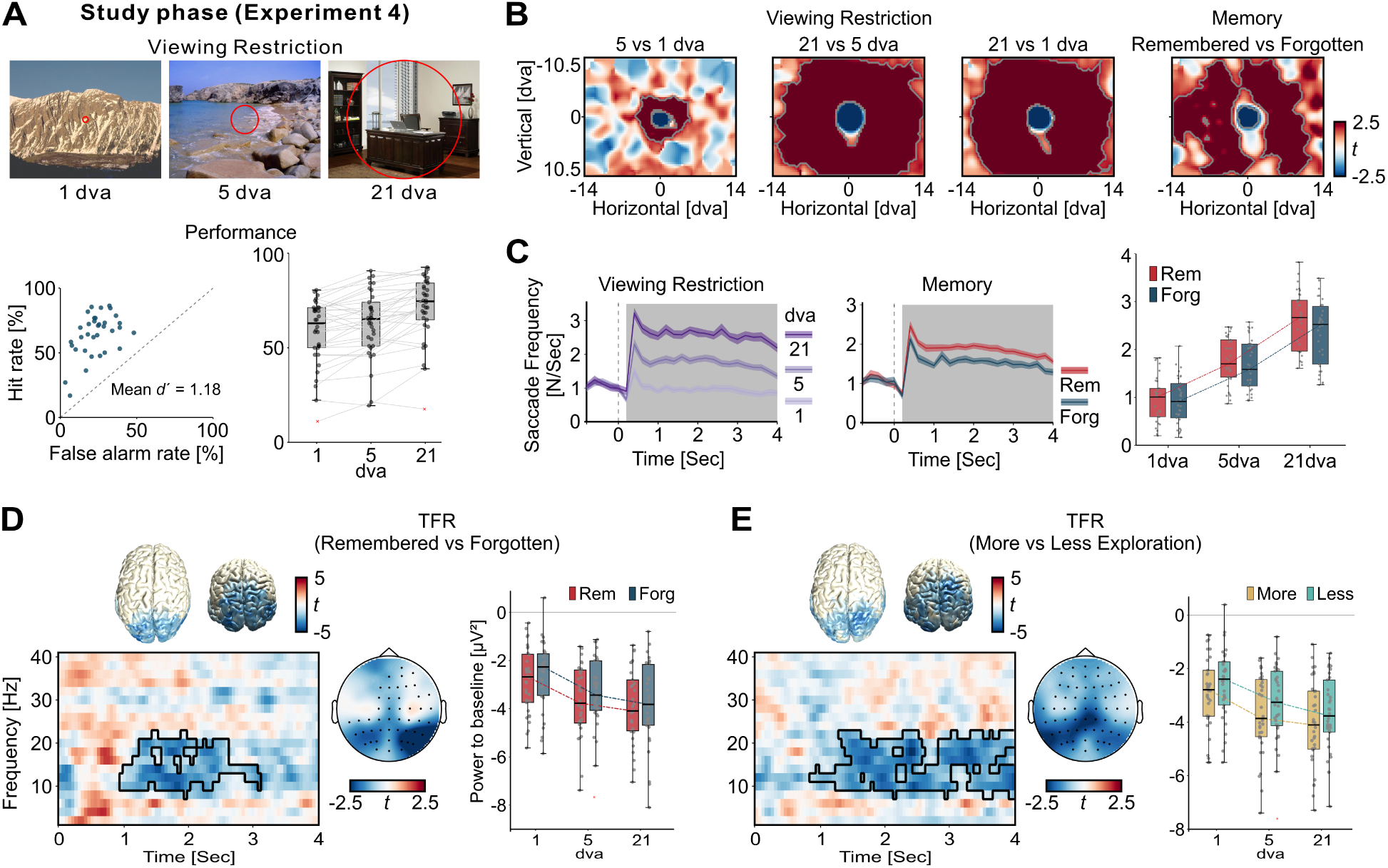
Restricting visual exploration modulates the degree of alpha/beta power reduction. (**A**) Experimental paradigm for Experiment 4 (top) and the memory performance (bottom). At the study phase, participants were instructed to explore the scenes only within a red circle (1/5/21 dva in diameter, shown at actual scale relative to the scene) displayed at the centre of the scene throughout the 4-sec viewing period. Trial structure at the study phase and subsequent test phase are identical to Experiment 1, 2 & 3. Memory performance across participants was above chance (bottom left), yet varied significantly as a function of viewing restriction (*F*(2, 66) = 37.57, *p* < 0.001; bottom right). (**B**) Gaze density contrasts revealed a bias towards surrounding scene locations and less towards the centre when visual exploration was less restricted (i.e., larger circle size). A similar bias was observed for later remembered versus forgotten scenes (right). (**C**) Greater eye movements at the study phase were observed when viewing scenes under less restrictive conditions (left), as well as for later remembered compared to forgotten scenes (middle). Average saccade frequency (right) showed significant main effects of both viewing restriction and memory. (**D**) An alpha/beta power subsequent memory effect was found, with greater reductions centred over posterior brain regions during scene viewing (left). Average power within the range of the effect (1–3 sec, 10–20 Hz, posterior electrodes) showed significant main effects of both viewing restriction and memory (right). (**E**) Across all levels of viewing restriction, more visually explored scenes were associated with greater reductions in alpha/beta power over posterior regions. Average power within the range of the effect (1–4 sec, 10–20 Hz, posterior electrodes) showed significant main effects of both viewing restriction and degree of exploration. Less restrictive viewing led to greater alpha/beta power reductions, and within each restriction level, higher degrees of visual exploration further enhanced the reduction. (**B–E**) Significant clusters are indicated by grey regions and grey or black contours (cluster-based permutation test, two-tailed, *p* < 0.05). In all boxplots, the central mark is the median across participants and the edges of the box are the 25^th^ and 75^th^ percentiles. Grey dots represent individual data points.

#### Viewing restriction reduced visual exploration and memory performance

Overall, participants remembered 64.11% (SD = 16.44%) of the scenes (Mean *d’* = 1.18, SD = 0.42; Fig. 4A, bottom left). However, memory performance varied significantly as a function of viewing restriction. Scenes viewed under more restriction (e.g., 1 dva condition) were remembered significantly less accurately than those viewed under less restriction (e.g., 21 dva condition). On average, participants remembered 58.20% (SD = 17.29%) of scenes in the 1 dva condition, compared to 62.94% (SD = 17.41%) in the 5 dva condition, and 71.19% (SD = 16.95%) in the 21 dva condition. A repeated-measures ANOVA confirmed a significant main effect of viewing restriction on memory performance (Fig. 4A, bottom right; *F*_(2, 66)_ = 37.57, *p* < 0.001, all pairwise comparisons with *p* < 0.001).

We first checked whether the viewing restrictions effectively constrained participant’s visual exploration as intended. The results showed it was indeed the case: gaze density contrasts revealed a significantly stronger bias towards surrounding scenes locations and reduced central fixation when the viewing restriction was lower (Fig. 4B, left; all clusters with *p* < 0.05, two-tailed, corrected for multiple comparisons across space). Similarly, less viewing restriction also led to significantly higher saccade frequency during scene viewing (Fig. 4C, left, *p* < 0.05, *F*-test, corrected for multiple comparisons across time). When trials were separated based on subsequent memory performance, we found that later remembered scenes showed a stronger bias towards surrounding locations and less towards the centre (Fig. 4B, right; all clusters with *p* < 0.05, two-tailed, corrected for multiple comparisons across space), as well as higher saccade frequencies (Fig. 4C, middle, *p* < 0.05, two-tailed, corrected for multiple comparisons across time) as compared to later forgotten scenes. A 3 x 2 repeated-measures ANOVA on trial-averaged saccade frequency showed significant main effects of viewing restriction (*F*_(2,66)_ = 101.36, *p* < 0.001), memory performance (*F*_(1,33)_ = 12.02, *p* < 0.001), as well as their interaction (*F*_(2,66)_ = 10.46, *p* < 0.001), indicating that the relationship between saccade frequency and memory performance varied across levels of restriction (Fig. 4C, right). Pairwise comparisons revealed significantly higher saccade frequencies for later remembered scenes in the 21 dva condition (*p* < 0.001, corrected for multiple comparisons), but not in the 5 dva condition (*p* = 0.197) and the 1 dva condition (*p* = 0.463). The results observed under the higher levels of viewing restriction (i.e., 1 and 5 dva) are consistent with Experiment 3, in which no difference in saccade frequency was found under attempted fixation (i.e., on a 1 x 1 dva fixation cross).

#### Viewing restriction modulates the degree of alpha/beta power reduction

We observed an alpha/beta SME, with greater alpha/beta power reductions over posterior regions of the brain that were later remembered as compared to those that were forgotten (Fig. 4D, left; *p* < 0.05, two-tailed, corrected for multiple comparisons across time, frequency and sensors). To further assess the impact of viewing restriction on this effect, we averaged the power within the identified time-frequency-electrode window of the SME (1–3 sec, 10–20 Hz, posterior electrodes) (Fig. 4D, right). A repeated-measure ANOVA yield significant main effects of viewing restriction (*F*_(2,66)_ = 22.28, *p* < 0.001) and memory performance (*F*_(1,33)_ = 6.08, *p* = 0.019), but no significant interaction (*F*_(2,66)_ = 0.74, *p* = 0.48). This suggests that, while an overall alpha/beta SME was present, the extent of alpha/beta power reduction was strongly modulated by the level of viewing restriction—less restrictive conditions (e.g., 21 dva) were associated with stronger alpha/beta power reductions.

We next examined whether the link between visual exploration and alpha/beta power persists across different levels of viewing restriction. Given that visual exploration was directly impacted by levels of viewing restriction (Fig. 4B), we median-split the trials based on their exploration index within each viewing restriction condition and averaged across conditions. The results confirmed that more visually explored scenes elicited greater alpha/beta power reductions, particularly over the posterior brain regions (Fig. 4E, left; *p* < 0.05, two-tailed, corrected for multiple comparisons across time, frequency and sensors). To further examine how alpha/beta modulation was jointly affected by viewing restriction and visual exploration, we averaged the power within the range of the detected effect (1–4 sec, 10–20 Hz, posterior electrodes) (Fig. 4E, right). A 3 x 2 repeated-measures ANOVA yielded significant main effects of viewing restriction (*F*_(2,66)_ = 25.91, *p* < 0.001) and visual exploration (*F*_(1,33)_ = 14.63, *p* < 0.001), but no significant interaction (*F*_(2,66)_ = 0.10, *p* = 0.908). Pairwise comparisons showed that within each level of viewing restriction, more visually explored scenes consistently elicited greater alpha/beta power reductions compared to less explored scenes (all pairwise comparisons with *p* < 0.05).

In sum, the results from Experiment 4 demonstrate that greater imposed viewing restrictions lead to poorer memory performance and also attenuate overall alpha/beta power reductions, irrespective of memory performance. However, within each restriction condition, scenes associated with greater visual exploration consistently showed stronger alpha/beta power reductions. Together, these findings suggest that visual exploration plays a key role in modulating alpha/beta activity during scene viewing: greater visual exploration, either achieved by less externally imposed restrictions on viewing or driven by more self-exploratory behaviour, leads to greater alpha/beta power reduction.

### 2.5. Saccades’ size predicts posterior alpha/beta amplitude

Because of the consistent association between the degree of visual exploration and alpha/beta power reduction under different viewing settings, we next sought out to explore whether eye movements themselves underlie this neural modulation. If that is indeed the case, we would expect to observe a direct link between saccade parameters and alpha/beta amplitude. Based on our results from scalp and intracranial EEG, we filtered the data in the 10–20 Hz range using a Hilbert transform to extract the amplitude envelop of alpha/beta activity. We then aligned scalp and intracranial EEG data to the onsets of detected saccades and extracted ±0.5 sec of EEG data around each onset. For scalp EEG, we focused on nineteen posterior electrodes to delineate the effect observed across the experiments (see Methods for details). To evaluate whether and how alpha/beta amplitude is modulated by saccades, we first generated a shuffled control condition by randomly permuting the trial labels from which the saccades were drawn, repeating this procedure 100 times. The resulting ’false’ saccade-locked activity was averaged across these permutations and served as a baseline for comparison with the true saccade-locked signal. In addition, we sorted all saccades by size and divided them into three equal portions: small, medium, and large. The saccade-locked EEG activity was then averaged within each portion to assess how saccade magnitude modulates alpha/beta activity.

During free scene viewing in Experiment 1, we found that saccades were associated with modulations in posterior alpha/beta activity, beginning approximately ∼0.1 sec prior to saccade onset and persisting until ∼0.35 sec after (Fig. 5A, bottom left). Specifically, saccades induced a brief decrease in alpha/beta amplitude preceding their onset, followed by an increase peaking at ∼ 0.12 sec post-onset, then a subsequent decline for the next ∼0.1 sec and an eventual return to baseline. Importantly, saccade size predicted the dynamics of post-saccadic alpha/beta amplitude. When we divided saccades based on their size (small: M = 1.67 dva, SD = 0.57; medium: M = 4.35 dva, SD = 1.27; large: M = 9.26 dva, SD = 1.47; Fig. 5A, top middle), we found that large saccades led to a significantly delayed peak (small: 0.123 sec, SD = 0.009; medium: M = 0.127 sec, SD = 0.008; large: M = 0.135 sec, SD = 0.009; *F*_(2,38)_ = 17.42, *p* < 0.001; all pairwise comparisons with *p* < 0.05) and a stronger peak alpha/beta amplitude (small: 0.240 mV, SD = 0.159; medium: M = 0.414 mV, SD = 0.230; large: M = 0.585 mV, SD = 0.301; *F*_(2,38)_ = 29.46, *p* < 0.001; all pairwise comparisons with *p* < 0.001) (Fig. 5A, top right). This saccade size-dependent modulation was clearly visible in the grand-averaged data (Fig. 5A, bottom right), where alpha/beta activity peaked later (small: 0.124 sec; medium: 0.128 sec; large: 0.136 sec) and higher (small: 0.232 mV; medium: 0.403 mV; large: 0.567 mV) with increasing saccade size. Notably, when comparing the post-saccadic alpha/beta amplitude for saccades made during the viewing of later remembered versus forgotten scenes, no significant differences were found in either peak amplitude (remembered: M = 0.390 mV, SD = 0.189; forgotten: M = 0.433 mV, SD = 0.268; *t*_(19)_ = -1.30, *p* = 0.209) or peak latency (remembered: M = 0.130 sec, SD = 0.008; forgotten: M = 0.129 sec, SD = 0.007; *t*_(19)_ = 1.01, *p* = 0.326) (Supplementary Fig. 5A).

**Figure 5.**
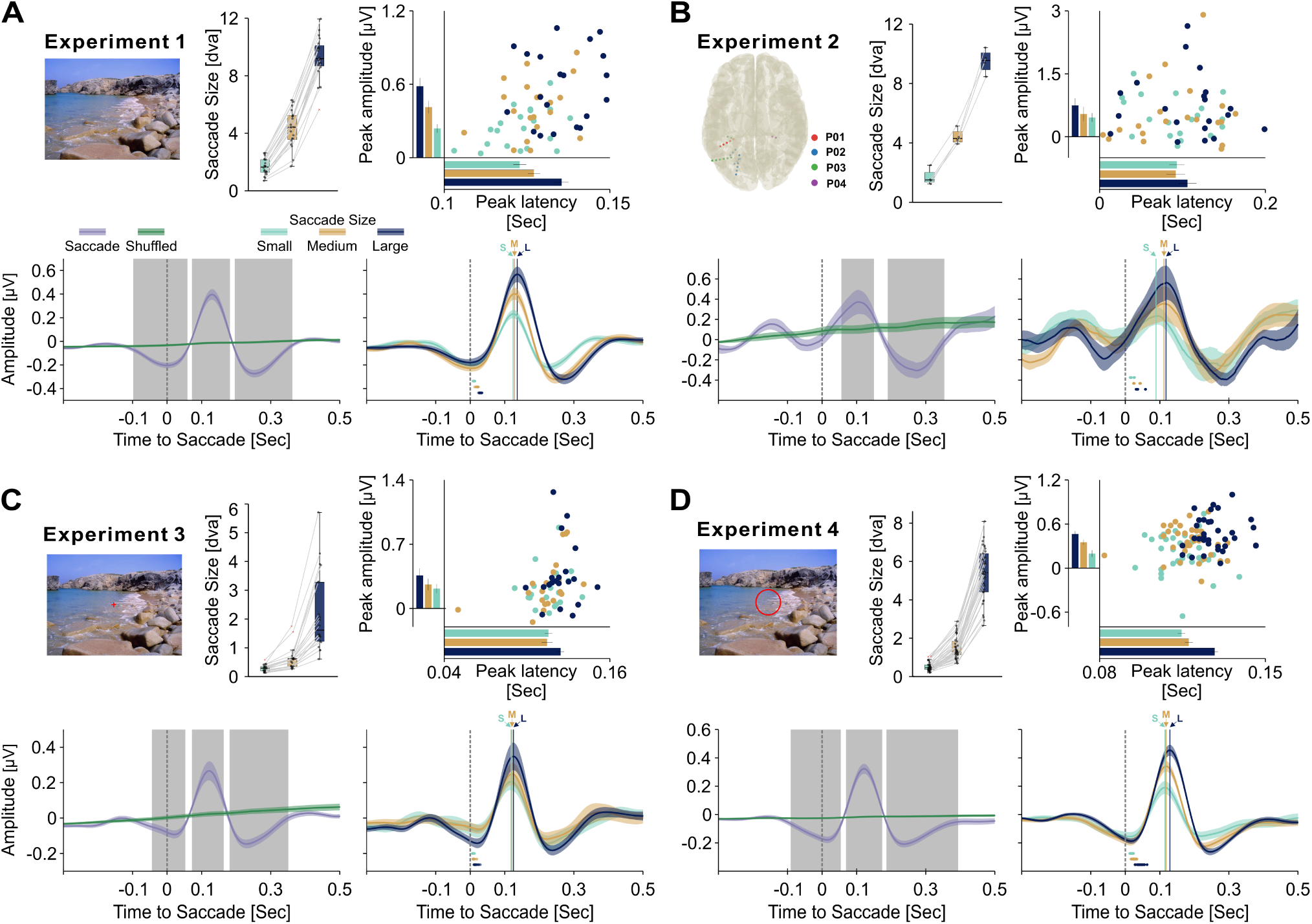
Saccades parameters predict posterior alpha/beta amplitude. (**A**) During free picture viewing, saccades were associated with modulations in posterior alpha/beta activity (bottom). Grey shading indicates time windows with significant differences between saccade-locked alpha/beta activity and shuffled data (cluster-based permutation test, *p* < 0.05, two-tailed). Additionally, when saccades were equally divided into small, medium, and large bins based on size (top middle), larger saccades were associated with a delayed yet stronger peak in alpha/beta amplitude (top right). Each dot represents one participant under the corresponding saccade condition. Group-level results (bottom right) confirmed this pattern: grand-averaged alpha/beta activity peaked later and at higher amplitude following larger saccades. (**B–D**) Similar patterns were observed across different datasets: (**B**) intracranial posterior electrodes during free viewing (Experiment 2), (**C**) posterior scalp electrodes during attempted fixation (Experiment 3), and (**D**) posterior scalp electrodes across varying degrees of viewing restriction (Experiment 4). In all boxplots, the central mark is the median across individual data points and the edges of the box are the 25^th^ and 75^th^ percentiles. Grey dots represent individual data points (bipolar-referenced channels in B; participants in all other panels). In each bottom right panel, small dots right to the vertical dashed line indicate the average offset time of individual participants’ saccades.

Following the same approach, we examined this pattern across different datasets. The clinical cohort with intracranial recordings also showed a modulation of posterior alpha/beta amplitude by saccade size (Fig. 5B, bottom left). Saccades were divided into three portions based on amplitude (small: M = 1.69 dva, SD = 0.56; medium: M = 4.42 dva, SD = 0.52; large: M = 9.50 dva, SD = 0.82). Across channels, there was a marginally significant effect of saccade size on peak alpha/beta amplitude (small: 0.459 mV, SD = 0.477; medium: M = 0.543 mV, SD = 0.761; large: M = 0.747 mV, SD = 0.743; *F*_(2,38)_ = 3.12, *p* = 0.056), though no significant effect on latency was found (small: 0.093 sec, SD = 0.041; medium: M = 0.092 sec, SD = 0.050; large: M = 0.106 sec, SD = 0.049; *F*_(2,38)_ = 0.79, *p* = 0.461), likely due to high variability across channels. Nevertheless, group-level results revealed a consistent trend: grand-averaged alpha/beta activity peaked later (small: 0.089 sec; medium: 0.112 sec; large: 0.118 sec) and at higher amplitude (small: 0.235 mV; medium: 0.355 mV; large: 0.564 mV) as saccade size increased. Further, comparisons based on memory performance again showed no significant differences in either peak amplitude (remembered: M = 0.483 mV, SD = 0.545; forgotten: M = 0.627 mV, SD = 0.616; *t*_(19)_ = -1.64, *p* = 0.117) or peak latency (remembered: M = 0.111 sec, SD = 0.039; forgotten: M = 0.100 sec, SD = 0.048; *t*_(19)_ = 0.98, *p* = 0.339) (Supplementary Fig. 5B).

Notably, in Experiment 3, although a large proportion of saccades consisted of microsaccades under attempted fixation (small: M = 0.29 dva, SD = 0.12; medium: M = 0.63 dva, SD = 0.39; large: M = 2.19 dva, SD = 1.39), the same trend was observed. Across participants, there was a marginally significant effect of saccade size on both the peak amplitude of alpha/beta activity (small: 0.217 mV, SD = 0.215; medium: M = 0.262 mV, SD = 0.281; large: M = 0.360 mV, SD = 0.348; *F*_(2,38)_ = 2.96, *p* = 0.064) and peak latency (small: 0.116 sec, SD = 0.012; medium: 0.115 sec, SD = 0.018; large: M = 0.124 sec, SD = 0.011; *F*_(2,38)_ = 3.09, *p* = 0.057). Group-level results were consistent, showing that grand-averaged alpha/beta activity peaked later (small: 0.118 sec; medium: 0.121 sec; large: 0.125 sec) and reached higher amplitude (small: 0.205 mV; medium: 0.250 mV; large: 0.348 mV) with increasing saccade size. Again, comparisons based on memory performance showed no significant differences in either peak amplitude (remembered: M = 0.269 mV, SD = 0.235; forgotten: M = 0.277 mV, SD = 0.266; *t*_(19)_ = -0.41, *p* = 0.684) or peak latency (remembered: M = 0.119 sec, SD = 0.008; forgotten: M = 0.120 sec, SD = 0.011; *t*_(19)_ = -0.39, *p* = 0.704) (Supplementary Fig. 5C).

In Experiment 4, we observed the same pattern. Saccade size (small: M = 0.51 dva, SD = 0.22; medium: M = 1.51 dva, SD = 0.58; large: M = 5.35 dva, SD = 1.35) predicted both peak latency (small: 0.115 sec, SD = 0.009; medium: 0.118 sec, SD = 0.010; large: M = 0.129 sec, SD = 0.008; *F*_(2,66)_ = 64.38, *p* < 0.001; all pairwise comparisons with *p* < 0.05) and peak amplitude (small: 0.197 mV, SD = 0.269; medium: M = 0.351 mV, SD = 0.215; large: M = 0.464 mV, SD = 0.228; *F*_(2,66)_ = 29.44, *p* < 0.001; all pairwise comparisons with *p* < 0.001). At the group level, larger saccades were consistently associated with a significantly delayed (small: 0.115 sec; medium: 0.118 sec; large: 0.129 sec), yet stronger (small: 0.190 mV; medium: 0.341 mV; large: 0.454 mV), peak in alpha/beta activity. When comparing based on memory performance, we found that saccades made during the viewing of scenes that were later remembered elicited a significantly higher peak alpha/beta amplitude (remembered: M = 0.347 mV, SD = 0.202; forgotten: M = 0.316 mV, SD = 0.218, *t*_(33)_ = 2.17, *p* = 0.037), although there was no significant difference in peak latency (remembered: M = 0.122 sec, SD = 0.010; forgotten: M = 0.121 sec, SD = 0.008, *t*_(33)_ = 0.51, *p* = 0.615) (Supplementary Fig. 5D). Note that this effect was driven by the tendency for saccades from remembered trials to originate from conditions with fewer viewing restrictions (on average, 52.22% (SD = 12.25%) of saccades from remembered trials occurred under the 21 dva condition, significantly higher than the ratio for forgotten trials: M = 39.72%, SD = 8.52%; *t*_(33)_ = 5.71, *p* < 0.001) and, consequently, with larger average saccade size (saccade size in remembered trials: M = 2.61 dva, SD = 0.67; forgotten trials: M = 2.15 dva, SD = 0.60; *t*_(33)_ = 6.27, *p* < 0.001). This interpretation aligns with the saccade size-dependent dynamics of post-saccadic alpha/beta amplitude consistently observed across experiments.

In sum, across all four independent datasets, saccades were consistently associated with modulations in posterior alpha/beta amplitude. Specifically, saccade size influenced the temporal dynamics, with larger saccades leading to delayed yet stronger peaks in alpha/beta amplitude. These results further clarify the functional relationship between visual exploration and neural oscillatory activity during scene viewing.

### 2.6. Alpha/beta amplitude reductions can be explained by accumulated eye movements

Finally, building on the finding that saccade size predicts alpha/beta amplitude, and the consistent observation that greater visual exploration is associated with stronger alpha/beta power reductions, we asked whether, more broadly, alpha/beta power reductions could be explained by the accumulation of eye movements over time.

To test this, we focused on the alpha/beta amplitude during the 1-sec interval following each saccade onset. We divided saccade-locked alpha/beta activity based on whether there was any subsequent saccade occurring within that 1-sec window (see Methods for details). Each saccade-locked activity was baseline-corrected to the post-saccadic interval of 0 to 0.2 sec and averaged separately for trials with as compared to without subsequent saccade.

During free scene viewing in Experiment 1, we found that post-saccadic alpha/beta amplitude was significantly reduced when followed by one or more saccades, compared to when no additional saccade occurred (Fig. 6A). While alpha/beta amplitude typically increased starting ∼0.25 sec after a saccade, subsequent saccades (most frequently occurred around 0.25 sec, as shown in the onset distribution in the lower panel) disrupted this rebound, leading to a lower amplitude across time. Importantly, this pattern was replicated across all datasets: in intracranial posterior electrodes during free viewing (Experiment 2; Fig. 6B), on posterior scalp electrodes under attempted fixation (Experiment 3; Fig. 6C), and under varying degrees of viewing restriction (Experiment 4; Fig. 6D).

**Figure 6.**
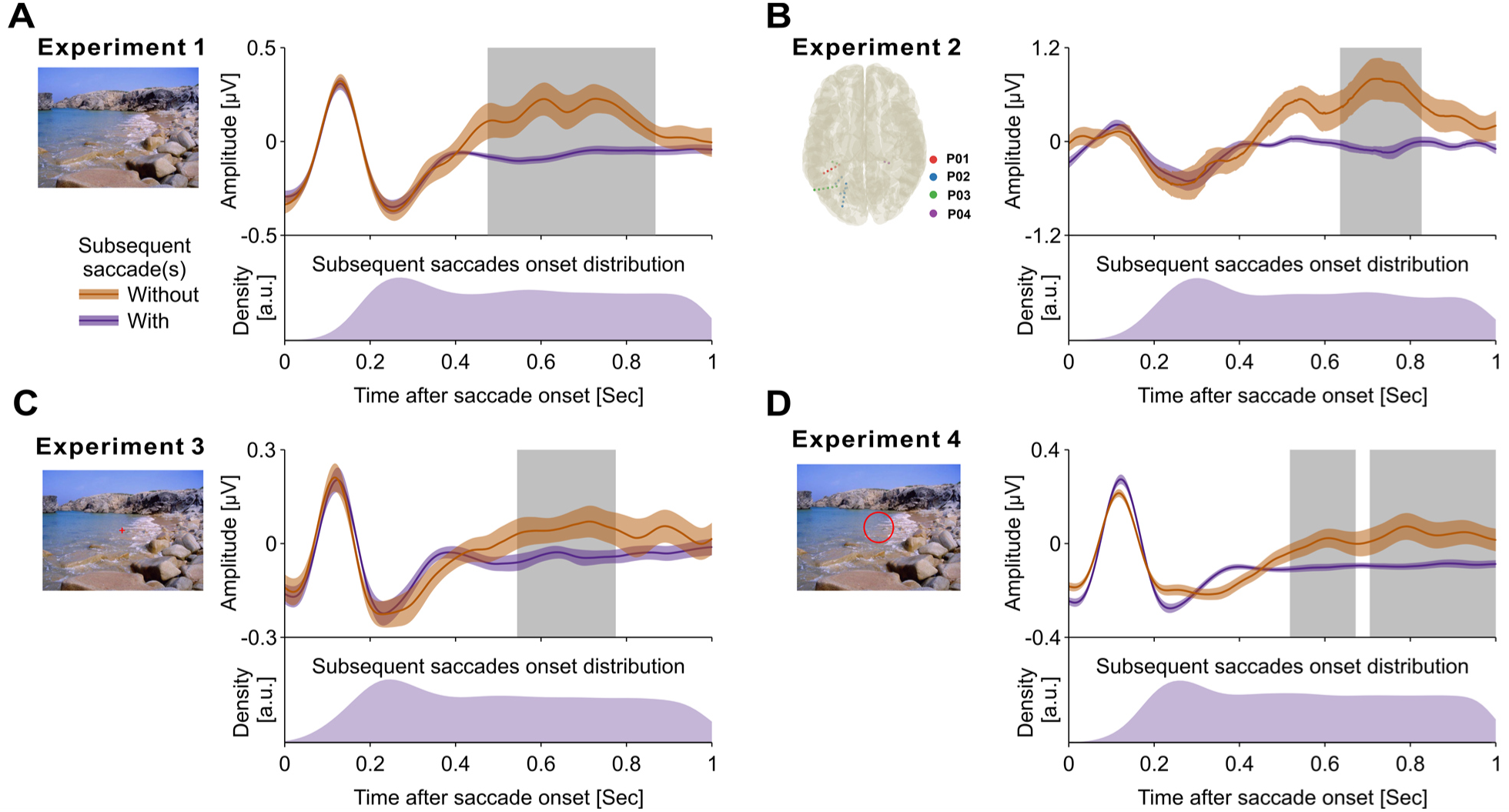
Alpha/beta amplitude reductions can be explained by accumulated eye movements. (**A**) Post-saccadic alpha/beta amplitude was significantly reduced when followed by additional saccades, compared to when no further saccades occurred. Grey shading marks time windows with significant clusters (cluster-based permutation test, *p* < 0.05, two-tailed). The distribution of subsequent saccades is shown below in arbitrary unit (a.u.). (**B–D**) Same patterns were observed in (**B**) intracranial posterior electrodes during free viewing, (**C**) posterior scalp electrodes during attempted fixation (Experiment 3), and (**D**) posterior scalp electrodes across varying degrees of viewing restriction (Experiment 4).

## 3. Discussion

The present results reveal a direct coupling between eye movements and alpha/beta activity that provides an alternative view on the correlation between alpha/beta desynchronization and subsequent memory performance, the so-called alpha/beta subsequent memory effect (SME). First, we showed that visual exploration and alpha/beta activity covary at the study phase to jointly predict memory performance at test, in line with recent work^28^. Going beyond previous work, we demonstrate this covariation in intracranial electrophysiology of posterior brain regions. To directly test the dependence of alpha/beta activity on eye movements, we manipulated participants’ gaze behavior. Importantly, when differences in visual exploration between later remembered and forgotten scenes were minimized by drastically restricting eye movements, the alpha/beta SME was absent. However, differences in alpha/beta power always emerged as soon as conditions with more eye movements were compared to conditions with fewer eye movements. Furthermore, the level of eye movement restriction impacted memory, with more restrictions resulting in worse memory performance. We also found that saccade size modulated alpha/beta amplitude across multiple viewing conditions, and, importantly, that reductions in alpha/beta power can be explained by the accumulation of saccades over time. Together, these findings advocate for a new perspective on the functional implications of alpha/beta neural activity and their relation to episodic memory formation, highlighting a tight coupling to eye movements.

The alpha/beta SME identified in Experiments 1, 2 & 4 is in line with previous studies, where greater reductions in alpha/beta power are observed for later remembered as compared to forgotten materials^6,8,30,40–43^. Although all of these studies presented visual stimuli that triggered visual exploration by the participants, data of how participants explored the stimuli (e.g., via eye tracking) is usually not available. Our work demonstrates that neglecting eye movements results in an incomplete understanding of the alpha/beta SME. Across experiments, we consistently found that a greater degree of visual exploration, quantified either by eye movement counts or spatial gaze distribution, was associated with stronger reductions in alpha/beta power. These reductions matched the frequency and spatial patterns typically attributed to the alpha/beta SME^6,8,30^. It is important to note that visual exploration itself strongly correlates with subsequent memory performance, and our data indicates that this correlation underlies the alpha/beta SMEs. In other words, both our current and other previous studies show that later remembered stimuli show higher levels of visual exploration during study^4,16–19^. This increased exploration is reflected in stronger reductions in alpha/beta power. Consequently, when compared to later forgotten stimuli, which are associated with less visual exploration at study, this contrast gives rise to the alpha/beta SME.

Another reason why visual exploration should be considered when interpreting the alpha/beta SME is the inevitability of eye movements and their direct impact on neural activity. Electrophysiological studies commonly instruct the participants to attentively fixate during the study phase to minimize movement-related artifacts in the signals (see e.g.,^44^). However, it is important to note that a presumed voluntary control over eye movements and the instruction to fixate does not result in completely stillness of the eyes^24,39^. Contrarily, eye movements of some form, large or small, are indispensable for accurate visual perception^45^. In the current study, we show that instructing participants to restrict their eye movements does not abolish eye movements completely, in particular when taking fixational eye movements such as microsaccades into account. More importantly, we found that also when fixation is attempted, visual exploration is directly coupled with a modulation of alpha/beta power. Across multiple data sets, cohorts and data types (EEG, iEEG), this link consistently shows, even if visual exploration is restricted experimentally. Together, these results suggest a strong coupling of eye movements and brain activity in the alpha/beta frequency range, and undermine an interpretation of the alpha/beta decrease as being predominantly driven by memory processes.

While this paper focusses on alpha/beta activity, it would be interesting to investigate other neural predictors of memory performance regarding their relationship with eye movements^2,3^. Theta activity, for example, has consistently been shown to correlate with subsequent memory performance^46^. Several studies also point at a direct link of theta activity and eye movements, in particular in the hippocampus of humans and non-human primates^47–52^, but also in other areas of the limbic circuit^53^. Regarding memory processes, previous studies either indicated a direct impact of eye movements on the hippocampal electrophysiology that predicts memory^47^ or an active role of hippocampal theta activity in coordinating eye movements^51,52^. Despite different functional interpretations, the commonality between our current and previous studies is that eye movements are mandatory to understand the link between brain activity and memory processes (see^54^).

To further substantiate the relationship between eye movements and alpha/beta activity, we asked whether eye movement parameters predicted the characteristics of alpha/beta activity. By investigating electrophysiology time-locked to individual saccades, we show that saccade size predicts the amplitude of alpha/beta activity, with larger saccades being associated with a larger post-saccadic peak in alpha/beta amplitude. Moreover, the post-saccadic alpha/beta activity peaked later as saccade size increased. Our findings thus add detailed information from saccade-locked analyses to previous observations which have described a modulation of neural activity through eye movements on a more global level^26,36,55,56^. For instance, the direction of saccades has been associated with transient lateralization of alpha activity^23,26,27^. Additionally, saccade size has been shown to modulate EEG amplitude both prior to and following saccade onset^57,58^. Importantly, our findings from multiple data sets including both scalp and intracranial recordings reveal that saccade size, but not subsequent memory performance, predicts the dynamics of alpha/beta neural activity. This dissociation further underscores eye movements as the more direct modulator of alpha/beta activity than memory. However, it is important to emphasize that our results do not argue against the functional coupling of eye movements and neural activity in service of memory formation. In fact, saccades have been shown to modulate alpha/beta neural activity in memory-relevant areas^48,49,59–61^, presumably to facilitate successful memory formation^59,60,62^. Our findings on saccade-onset locked alpha/beta activity are also in line with previous work on a fixation-related potential, the so-called lambda wave^63^. Similar to our findings, lambda waves exhibit a peak latency of around 100 milliseconds after fixation, and this latency varies with saccade size^36^. That the presence of lambda waves and alpha oscillations are correlated^64^ and the lambda wave’s spectral content is in the alpha range^36^ might indicate that both reflect the same underlying brain activity, a notion that has been debated extensively (see, e.g.,^65–67)^.

While investigating the consequences of single eye movement clarifies their immediate impact on electrophysiology, it is also important to understand how this effect accumulates over time to explain the typical pattern described as alpha/beta SME, i.e., a general decrease in alpha/beta power after stimulus onset that provides the basis of a greater decreases for later remembered (with more eye movements) than forgotten items (with fewer eye movements). To investigate this, we split the data in saccade-locked segments with and without subsequent saccades. The results showed higher, sustained alpha/beta amplitudes for segments without subsequent saccades than with subsequent saccades. This indicates that accumulating saccades over time results in gradually suppressed alpha/beta activity. Note that alpha/beta SMEs are usually computed relative to a baseline. This baseline is typically a data segment where participants were instructed to not move their eyes and fixate on a fixation cross. According to the evidence presented here, such a baseline will have relatively high alpha power^24^, in particular when compared to other segments that involve visual stimulation, and their difference would result in the commonly observed decrease in alpha/beta power.

An important question that follows from our results is what function the coupling between visual exploration and alpha/beta activity might have. We propose that the role of alpha/beta neural activity lies in the control and coordination of eye movements. Recent evidence suggests alpha power reduction follows moment-to-moment modulations of eye movements^23^, and such coupling is preserved even when no visual information is required to be processed (e.g., in full darkness)^25^. Moreover, saccades have been shown to couple with phase of alpha oscillation across various viewing tasks^23,59,60^, indicating a direct involvement of alpha oscillations in controlling saccade coordination and execution. Our results support this interpretation by showing that alpha/beta amplitude is tightly coupled to both the timing and parameters of individual saccades during visual exploration. Specifically, we observed a peri-saccadic decrease in alpha/beta amplitude, followed by a sharp increase during fixation that drops approximately 120 milliseconds later, aligning well with (i) the typical saccadic inhibition interval^68–71^ and (ii) the minimal duration of fixations of around 100-150 milliseconds^72,73^.

Therefore, the here described decreases and increases in alpha/beta amplitude around saccade onset may reflect a duty-cycle for executing eye movements, with high amplitudes imposing inhibition on initiating eye movements, and low amplitudes reflecting a release of this inhibition^23^. Put differently, the function of high alpha/beta activity could be viewed as stabilizing fixation on a target. Along these lines, the here reported scaling of the alpha/beta amplitude with saccade size could be interpreted as reflecting the effort of the oculomotor system to impose fixation on ballistically moving eyes. Since larger eye movements are also faster than small eye movements (see main sequence), their kinematics might ask for a stronger effort when stabilizing the fixation after the movement. This oculomotor account of alpha/beta activity in posterior brain regions aligns with earlier work in humans^23,74^, and also monkeys, where electrical stimulation of the visual cortex elicits eye movements^75^. Thus, the function of alpha/beta activity could be that it blocks the initiation of eye movements to extrafoveal targets by increasing alpha amplitudes in the respective, retinotopic brain areas.

In conclusion, the present report brings together two robust predictors of memory, visual exploration and neural activity, and advocates for a new perspective on the alpha/beta SME by taking eye movements into account. The presented findings demonstrate that visual exploration is tightly coupled with alpha/beta neural activity and that neglecting eye movements during the study phase of a memory experiment could lead to incomplete conclusions about what predicts successful memory formation. On a broader perspective, the present data supports the view that alpha/beta activity reflects the dynamics of the oculomotor system. Consequently, it is this relationship that in turn effects the manifestation of higher-level cognitive processes.

## 4. Methods

### 4.1. Participants

A total of 26 and 22 participants were initially recruited for Experiments 1 and 3, respectively. Based on the exclusion criteria pre-registered for the studies (Experiment 1: https://osf.io/rhn4z/; Experiment 3: https://osf.io/czjeh/)—specifically, participants who either failed to complete the experiment or had fewer than 20 remembered or forgotten trials in the free-viewing memory task—a final sample size of N = 20 was included in each experiment (Experiment 1: 8 females; age range: 18–32 years; M = 23.77; Experiment 3: 13 females; age range: 19–33 years; M = 25.05). Sample size estimation was conducted using G*Power^76^, following the parameters reported in Popov & Staudigl (2023)^28^ for matched-samples t-tests (N = 16, α = 0.05, power = 0.95, *dz* ≈ 1). We opted for N = 20 in both experiments to account for potential overestimation of effect sizes. For Experiment 4, 36 participants were initially recruited. After excluding 2 participants who did not complete the task, the final sample included N = 34 participants (19 females; age range: 21–35 years; M = 25.97).

Participants received either monetary compensation or course credit for their participation. Recruitment was conducted through advertisements posted across the university and online announcements. All participants self-reported having no uncorrectable visual impairments, psychiatric or neurological disorders, (suspected) pregnancy, recent alcohol consumption (more than two alcoholic drinks within 24 hours prior to the experiment), history of alcohol abuse, ongoing or acute drug use, or use of prescription medications that may alter brain function. Additionally, for Experiment 3 and 4, participants were screened for colour vision deficiencies. Those who exhibited colour vision deficiency were excluded from further participation. The study was approved by the Ethics Committee of the Department of Psychology at Ludwig-Maximilians-Universität München. All participants provided informed consent prior to participation.

For the clinical cohort, four patients from the Epilepsy Centre, Department of Neurology at Ludwig-Maximilians-Universität München, all diagnosed with medically intractable epilepsy, volunteered to participate. This study was approved by the Ethics Committee of the Medical Faculty of Ludwig-Maximilians-Universität München, and all patients provided informed consent prior to participation.

### 4.2. Task design and procedure

#### Experiment 1

Each scene (size: 28 x 21 degrees of visual angle (dva); duration: 4 sec) was centrally displayed following a white central fixation cross (size: 1 x 1 dva; duration: 1.5 sec ± 0.2 sec jitter) (Fig.1A, left). Participants were instructed to freely view the scenes and subsequently indicate whether the scene depicted an indoor or outdoor environment via a button press. The next trial started immediately after the response. A total of 216 scene-viewing trials were presented (randomly drawn from a set of 324 scenes), divided into six blocks. Each block consisted of three parts, each comprising 12 consecutive trials, interleaved with a 2-minute visual detection task. This task is not described further as it is unrelated to the research questions and hypotheses of the current study. Participants were informed about the subsequent test phase prior to the start of the experiment.

Following the study phase, participants took a 10-minute break filled with distractor tasks (e.g., handedness and dominant eye assessments). The test phase then began (Fig. 1A, middle). Each scene (size: 28 x 21 dva; duration: 4 sec) was centrally displayed following a white central fixation cross (size: 1 x 1 dva; duration: 1.5 sec ± 0.2 sec jitter). Participants were instructed to freely view the scenes and then indicate whether the scene had been shown previously during the study phase (“old”) or not (“new”), and to rate their confidence in this judgment on a scale from 1 (“very unsure”) to 4 (“very sure”). Both the old/new recognition and confidence ratings were self-paced. The next trial began immediately after the confidence rating. The test phase comprised a total of 324 trials: 216 “old” scenes and 108 “new” scenes, equally split between indoor and outdoor categories.

#### Experiment 2

The clinical cohort followed the same overall trial structure as Experiment 1, with an adapted number of stimuli and block design to reduce task duration for patient comfort. The task was divided into two sessions. In each session, patients completed six blocks of 12 scene-viewing trials, interleaved with a 1-minute visual detection task. The test phase followed immediately after the study phase and included 108 trials per session: 72 “old” and 36 “new” scenes, evenly divided between indoor and outdoor categories. Across both sessions, this resulted in 144 scene-viewing trials and 216 test trials.

### Experiment 3

Each scene (size: 28 x 21 dva; duration: 4 sec) was centrally displayed following a red central fixation cross (size: 1 x 1 dva; duration: 1.5 sec ± 0.2 sec jitter) (Fig.3A, left). The central fixation cross remained visible on top of the scene, and participants were instructed to maintain fixation throughout the scene presentation. The total number of stimuli and block structure was the same as in Experiment 1. Each part of 12 consecutive trials was interleaved with two eye movement tasks (∼2 minutes in total). These tasks are not described further, as they are unrelated to the present research questions and hypotheses. As in Experiment 1, participants were informed about the test phase before the experiment began.

After the study phase and the 10-minute distractor task break, the test phase proceeded identically to that in Experiment 1.

#### Experiment 4

Each scene (size: 28 x 21 dva; duration: 4 sec) was centrally displayed following a red central fixation cross (size: 1 x 1 dva; duration: 1.5 sec ± 0.2 sec jitter) (Fig.4A, top). During scene presentation, a red circle was overlaid centrally on the display and remained visible throughout the 4-sec viewing period. Participants were instructed to view only the scene area within the boundary of the red circle. The experiment contained three viewing restriction conditions, defined by the diameter of the red circle (1, 5, or 21 dva). A total of 324 trials (randomly drawn from a set of 486 scenes) were divided into six blocks. Each block included all three viewing conditions, with 18 consecutive trials per condition. The order of conditions was counterbalanced within and across blocks.

Following the study phase and the 10-minute break filled with distractor tasks, participants proceeded to the test phase, which followed the same structure as in Experiments 1 and 2. The test phase comprised 486 trials: 324 “old” and 162 “new” scenes, evenly distributed across indoor and outdoor categories.

### 4.3. Data acquisition

#### Gaze data acquisition

Binocular gaze positions were tracked throughout all experiments using an eye tracker. Experiments 1 and 3 were recorded with the Tobii Pro Spectrum (sampling rate: 600 Hz), and Experiment 4 used the EyeLink (1000 Hz). For the clinical cohort in Experiment 2, gaze data were collected from two patients using the Tobii Pro Spectrum (600 Hz) and from two patients using the EyeLink (1000 Hz).

#### Scalp EEG acquisition

In Experiments 1, 3, and 4, scalp EEG was recorded using a 64-channel system (ANT Neuro) at a sampling rate of 1000 Hz. Electrode impedance was maintained below 10 kΩ. EEG signals were referenced online to electrode CPz. Additionally, horizontal and vertical electrooculogram (EOG) activity were simultaneously recorded.

#### Intracranial EEG acquisition

For Experiment 2 (clinical cohort), intracranial EEG was recorded from Spencer depth electrodes (Ad-Tech Medical Instrument, Racine, WI, USA), each with 4–12 contacts spaced 5 mm apart. Data were acquired using XLTEK Neuroworks software (Natus Medical, San Carlos, CA, USA) with an XLTEK EMU128FS amplifier. Signals were referenced to a parietal electrode and sampled at 1000 Hz.

### 4.4. Data preprocessing and analysis

#### 4.4.1. Gaze data

To quantify gaze variability during the 4-sec scene viewing period, horizontal and vertical gaze positions were binned into a two-dimensional (2-D) array (80 x 60 bins) matching the size of the image (28 x 21 dva). For each trial, counts in 2-D bin were normalized to the total counts of gaze positions and then smoothed using a 2-D Gaussian filter (with width parameter sigma set to 2.5, same as outline in Popov & Staudigl (2023)^28^). Additionally, we calculated an exploration index for each trial, represented by the standard deviation of the normalized and smoothed 2-D gaze density array. As standard deviation measures the spread around the mean, the exploration index reflects the spread of gaze, quantifying overall visual exploration during the 4-sec scene presentation and allowing for comparisons across trials (see Supplementary Fig.4 for illustration).

For saccade detection, we calculated the 2-D velocity based on the horizontal and vertical gaze positions. Samples exceeding a velocity threshold (defined relative to the trial-based median velocity) and lasting longer than 12 milliseconds were classified as saccades^34^. To optimize detection sensitivity across different experimental conditions, the velocity threshold was scaled accordingly: 6× the median velocity in Experiment 1 (free viewing), 4× for the clinical cohort and Experiment 4 (noisier settings, restricted viewing, respectively), and 3× in Experiment 3 to enhance detection of microsaccades (e.g., with saccade size < 1 dva). To exclude artifacts, saccades occurring within 100 milliseconds before or after a blink were automatically removed, and remaining false positives (e.g., abrupt changes in gaze position due to noise) were excluded via manual trial-by-trial inspection. Final saccades were defined as either (i) binocular saccades detected in both eyes with overlapping time windows, merged by taking the earliest onset and latest offset across eyes, or (ii) monocular saccades detected in only one eye when no corresponding event was found in the other (e.g., due to missing or noisy gaze data). Additionally, to prevent double-counting, a minimum inter-saccadic interval of 50 milliseconds was applied: any subsequent saccades occurring within the 50 milliseconds of a preceding saccade were discarded.

Saccade metrics-size, duration and direction-were calculated based on the difference in gaze position between saccade onset and offset. For binocular saccades, gaze positions from both eyes were averaged; for monocular saccades, calculations were based on the gaze data from the eye in which the saccade was detected.

#### 4.4.2. EEG data

##### Scalp EEG

EEG data analyses were performed using the open-source software FieldTrip^77^. Data were offline re-referenced to the average of all scalp electrodes. The continuous data were segmented and time-locked to scene onsets. Oculo-muscular and cardiac artifacts were detected and removed by means of independent component analysis (ICA) and noisy trials were excluded based on visual inspection.

##### Intracranial EEG

A total of 27 electrodes from four patients, all implanted in posterior brain regions, were included in the current study (for detailed electrode localisation, see Supplementary Table. 1). For each patient, electrode locations in the brain were determined by examining computed tomography (CT) and preoperative T1-weighted Magnetic Resonance Imaging (MRI) scans. Cerebral atlases were generated by parcellating the preoperative T1 images using FreeSurfer^78^. The CT scan was co-registered to the corresponding MRI, and electrode positions along with anatomical labels were extracted using YAEL toolbox^79^. A notch filter was used to suppress power at the electrical power line frequency of 50 Hz. All intracranial contacts were re-referenced in a bipolar montage using neighbouring contacts, resulting in N = 20 bipolar channels.

Interictal epileptic discharges (IEDs) were detected in each bipolar channel using an automated algorithm based on the approach described in Vaz et al. (2019)^80^. First, data were segmented to include only the relevant task period. For each time point, we calculated a z-score based on both the signal gradient (first derivative) and amplitude after applying a 250 Hz high-pass filter. Time points exceeding a z-score threshold of 5 in either the gradient or high-pass-filtered amplitude were identified. A window of ±0.25 sec surrounding each identified time point was marked as an IED event. Additionally, the data were epoched time-locked to image onset, and manually inspected to identify and mark any remaining epileptiform activity not detected by the automated procedure. Finally, we replaced all the marked IED events with NaNs for further analyses.

For both scalp and intracranial EEG, TFRs were computed using a sliding window of 0.5 sec and a Hanning taper, resulting in a frequency resolution of 2Hz (ranging from 2–40 Hz). Power estimates were averaged across trials for each condition (e.g., remembered vs forgotten) and baseline corrected to the time window of -1 to -0.5 sec before image onset.

#### 4.4.3. Source analysis

We applied a frequency-domain beamforming technique for source reconstruction of the scalp EEG^81^. This approach involves constructing a cross-spectral density (CSD) matrix for the frequency band of interest (e.g., 10–20 Hz, covering the alpha/beta range). Based on this matrix, spatial filters were generated for each voxel (1 cm³ resolution) to estimate source power at specific brain locations. These filters were computed using both data during the pre and post stimulus intervals (e.g., common filter approach). Anatomically head models were generated using standard MRI template provided by FieldTrip^77^. To improve visualization, we plotted the outcome of a cluster-based permutation statistics (using a *p* < 0.05, two-tailed, corrected for multiple comparisons across time, frequency and virtual sensors) in the source plot. Note, however, that the statistical inference whether there is a significant difference between conditions (e.g., later remembered versus later forgotten) was already derived from scalp-level data, making source statistics redundant.

#### 4.4.4. Saccade locked analysis

To further evaluate alpha/beta activity modulated by saccades, we aligned scalp and intracranial EEG data to the onset of detected saccades. For scalp EEG, we focused on nineteen posterior electrodes (Pz, P1/2, P3/4, P5/6, P7/8, POz, PO3/4, PO5/6, PO7/8, Oz, O1/2). This electrode selection was based on the following rationale: (i) the alpha/beta power modulations identified across Experiment 1, 3 & 4 were consistently source localised at posterior regions of the brain; (ii) these electrodes cover the spatial distribution of effects reported in prior studies characterising EEG activity modulated by eye movements ^23,25,27,28,38,56^.

For both scalp and intracranial EEG, the continuous data time-locked to image onset were filtered in the 10–20 Hz range using a Hilbert transform to extract the amplitude envelope of alpha/beta activity. EEG segments were then extracted ±0.5 sec around each detected saccade onset. Only saccades occurring at least 0.5 sec after image onset and before the final 0.5 sec of the 4-sec image presentation were included, ensuring that all extracted segments remained within the stimulus presentation time window. For scalp EEG, the filtered data were averaged across the selected posterior electrodes. Each saccade-locked segment was baseline-corrected using the pre-saccadic interval from -0.5 to -0.3 sec. To statistically evaluate saccade-related modulations in alpha/beta amplitude, a shuffled control condition was generated by randomly permuting the trial labels from which saccades were drawn. This permutation was repeated 100 times, and saccade-locked alpha/beta amplitudes for these “false” onsets were averaged across repetitions. The resulting shuffled amplitudes was then compared to the real saccade-locked alpha/beta amplitude for statistical analysis.

We next examined whether saccade parameters or subsequent memory performance predict the dynamics of post-saccade alpha/beta amplitude. Saccade-locked alpha/beta activity was submitted into different conditions based on the saccade size (split equally into three portions) or the memory performance of the trial (remembered or forgotten) from which each saccade was extracted. For each participant (or each channel in Experiment 2, clinical cohort), saccade-locked alpha/beta activities were averaged within conditions.

The Matlab function *findpeaks* was used to identify the most prominent peak within the first 0.2 sec following saccade onset. Peak latency and amplitude were then statistically assessed at the subject level across the respective conditions.

To further explore whether alpha/beta amplitude reductions can be explained by accumulated saccades, we focused on the 1-sec post-saccadic time window. We compared post-saccadic amplitude modulation based on whether or not there was any subsequent saccades. We further excluded the saccades occurred at any time point during the last 1 sec before end of the image presentation so that all the extracted neural activity fell within. The extracted alpha/beta activity is time-locked to the saccade onset and covered 1 sec after. Each trial was baseline corrected to the period of [0, 0.2] sec following saccade onset, and split and averaged within conditions based on whether there were any subsequent saccades in the next second, then for subsequent statistical testing.

#### 4.4.5. Statistics

To evaluate the difference in saccade frequency across time, gaze variability and neural activity between conditions, we used non-parametric cluster permutation tests with Monte Carlo randomization^82^. The approach identifies clusters across time points (or space points in case of gaze array), frequency, and electrodes (whenever applicable). The test employed a dependent samples *t*-test (or *F*-test when comparing more than two conditions) to generate the randomization distribution. The null hypothesis was tested by comparing the observed cluster-level statistics to the distribution obtained from permutations of condition labels. In the TFR analyses, we required a minimum of three neighbouring electrodes for cluster formation to increase spatial specificity.

To address the imbalance between remembered and forgotten trials observed in Experiment 1, 3 & 4, a trial-balancing procedure was applied. Specifically, for each participant, trials from the condition with more trials were randomly subsampled to match the number in the other condition. This was repeated 100 times, and the resulting gaze density arrays or TFRs were averaged across iterations prior to statistical testing.

## Acknowledgments

This work was supported by the Deutsche Forschungsgemeinschaft (DFG, German Research Foundation) – Project number 553351953 (awarded to X. Wu) and by the European Research Council (ERC, Starting Grant 802681 awarded to T. Staudigl). We are grateful to all participants, in particular to all patients, who volunteered to participate in this study. We thank the staff and physicians at the Epilepsy Center, Department of Neurology, Ludwig Maximilians University, Munich for assistance. We thank Soumya Kesavabhotla, Ritika Gupta, and Calvin Lange for their assistance with data collection.

## Author Contributions

X.W.: conceptualization, methodology, formal analysis, investigation, interpretation of results, data curation, writing — original draft, writing — review and editing, visualization, and funding acquisition. T.P.: methodology, and writing — review and editing. T.B.: investigation, writing — review and editing. N.F.: resources and writing— review and editing. C.V.: resources and writing — review and editing. E.K.: resources and writing — review and editing. J.R.: resources and writing — review and editing. T.S.: conceptualization, methodology, interpretation of results, data curation, writing — original draft, writing — review and editing, supervision, project administration, and funding acquisition.

## Competing Interests

The authors declare that they have no competing financial interests.

## Data availability

All datasets for healthy participants will be uploaded to a publicly available repository upon publication.

## Code availability

All code related to the analyses of the manuscript will be made publicly available upon publication.

## Supplementary Methods

### Memorability analysis

We quantified the visual memorability of the scene stimuli to examine the extent to which their intrinsic properties could account for differences in eye-movements across conditions. To this end, we used MemNet^29^, a pre-trained convolutional neural network (CNN) trained on 60,000 images, to estimate a memorability score for each image used in the current study. MemNet captures a range of image attributes such as saliency, popularity, and aesthetics etc., and returns a continuous score from 0 to 1 that reflects the objective likelihood of an image being remembered. This approach allowed us to test whether the intrinsic visual features of the stimuli could explain the observed variability in eye movements.

### Generalized linear mixed-effects model (GLMM)

To assess whether memory performance (binary: remembered = 1, forgotten = 0) was predicted by image memorability scores and saccade counts, we fitted a GLMM with a logit link function. In the first model, we included the *z*-transformed memorability score as a fixed effect and specified random intercepts for both subject and image (Supplementary Fig. 2A). We then extended this model by adding *z*-transformed saccade counts as an additional fixed effect, while maintaining the same random-effects structure (random intercepts for subject and image). Finally, we compared the two nested models using a likelihood ratio test to evaluate whether including saccade counts as a fixed effect significantly improved model fit.

### Linear mixed-effects model (LMM)

To assess whether the number of saccades made during viewing could be explained by intrinsic image properties, we fitted an LMM with *z*-transformed saccade count as the dependent variable and *z*-transformed image memorability score as a fixed effect. Random intercepts were included for both subject and image to account for within-subject and within-item variability (Supplementary Fig. 2B).

### Fitting Oscillations and One-Over-F (FOOOF) analysis

We applied the Fitting Oscillations and One-Over-F (FOOOF) algorithm to separate periodic oscillatory components from the aperiodic background. Specifically, we used the established method to parameterize the neural power spectral density into aperiodic (1/f-like) and periodic (oscillatory) components, extracting the oscillatory power above the fitted aperiodic background^31^. The algorithm was applied to trial-averaged TFRs following the trial-balancing procedure (where applicable), across all time points and channels. The resulting oscillatory power estimates were then baseline-corrected for each condition using absolute subtraction relative to the interval from -1 to -0.5 secs before image onset. Statistical comparisons between conditions were conducted using the same procedures as for the non-FOOOFed power estimates.

**Supplementary Figure 1.**
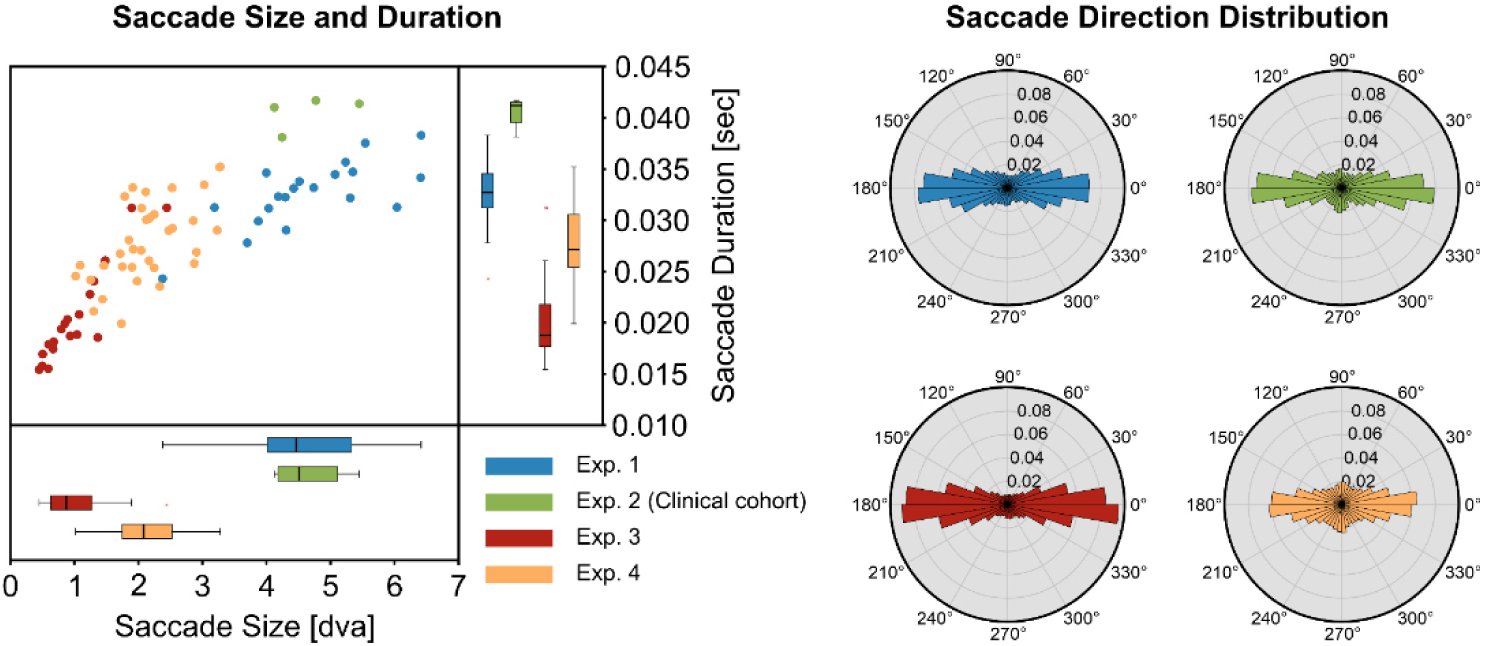
Saccade metrics across experiments. Subject-averaged saccade size and duration are shown on the left. Each dot represents an individual participant from the corresponding experiment. Saccade size was modulated by the task’s viewing instructions. Participants made significantly smaller saccades in Experiment 3 under attempted fixation (M = 1.00 dva, SD = 0.51), compared to varying degree of viewing restriction in Experiment 4 (M = 2.14 dva, SD = 0.61), and further compared to free viewing in Experiment 1 (M = 4.65 dva, SD = 1.04) and Experiment 2 (clinical cohort: M = 4.65 dva, SD = 0.60; 4-way ANOVA: *F*_(3,74)_ = 101.008, *p* < 0.001). Same pattern was found for saccade duration: participants made shorter saccades in Experiment 3 (M = 0.020 sec, SD = 0.005), compared to in Experiment 4 (M = 0.028 sec, SD = 0.004), and further compared to Experiment 1 (M = 0.033 sec, SD = 0.003) and Experiment 2 (clinical cohort: M = 0.041 sec SD = 0.002; 4-way ANOVA: *F*_(3,74)_ = 48.488, *p* < 0.001). However, across all experiments, saccade duration increased proportionally with saccade size. Subject-averaged saccade direction distributions are shown on the right.

**Supplementary Figure 2.**
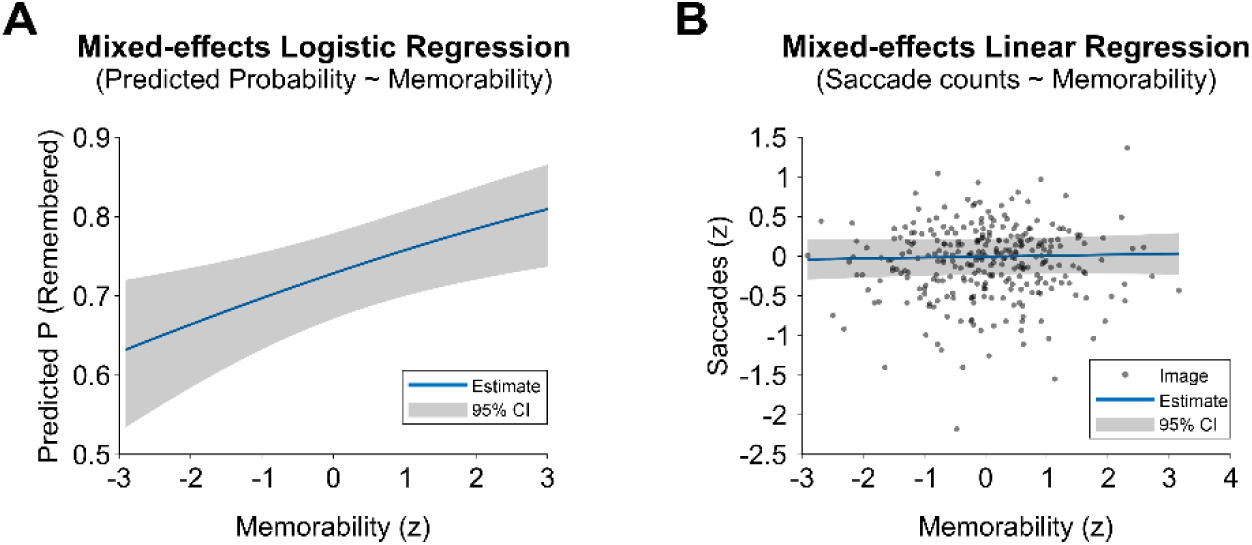
**Memorability of stimulus predicts memory performance but not saccade counts**. (**A**) Generalized linear mixed-effects model (GLMM) on memory performance (binary: remembered vs, forgotten) predicted by memorability of stimuli in Experiment 1. Higher memorability score of stimuli significantly predicted better memory performance (*p* = 0.004). (**B**) Linear mixed-effects model (LMM) on numbers of saccades conducted during viewing of novel scenes predicted by memorability of stimuli in Experiment 1. Image memorability did not significantly predict saccade counts during viewing (*p* = 0.58).

**Supplementary Figure 3.**
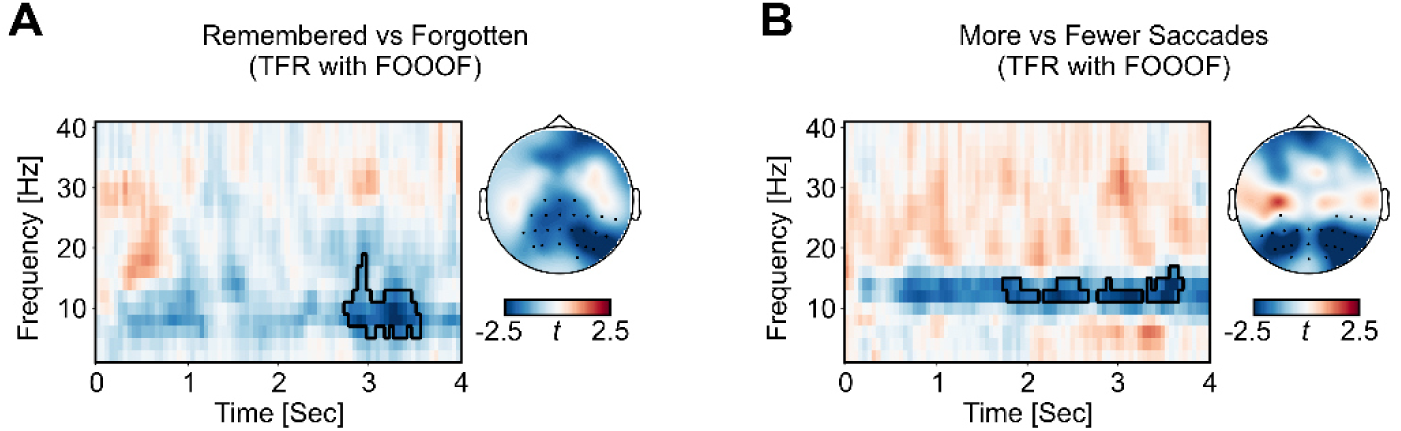
TFR contrasts using FOOOF-extracted periodic power. (**A**) Later remembered scenes elicited significantly greater oscillatory power reductions in the alpha/beta range compared to later forgotten scenes, showing a temporal and topographical distribution consistent with results based on non-FOOOFed power. (**B**) Trials viewed with more saccades showed greater oscillatory power reductions than those with fewer saccades, consistent with the temporal and topographical distribution of the results based on non-FOOOFed power.

**Supplementary Figure 4.**
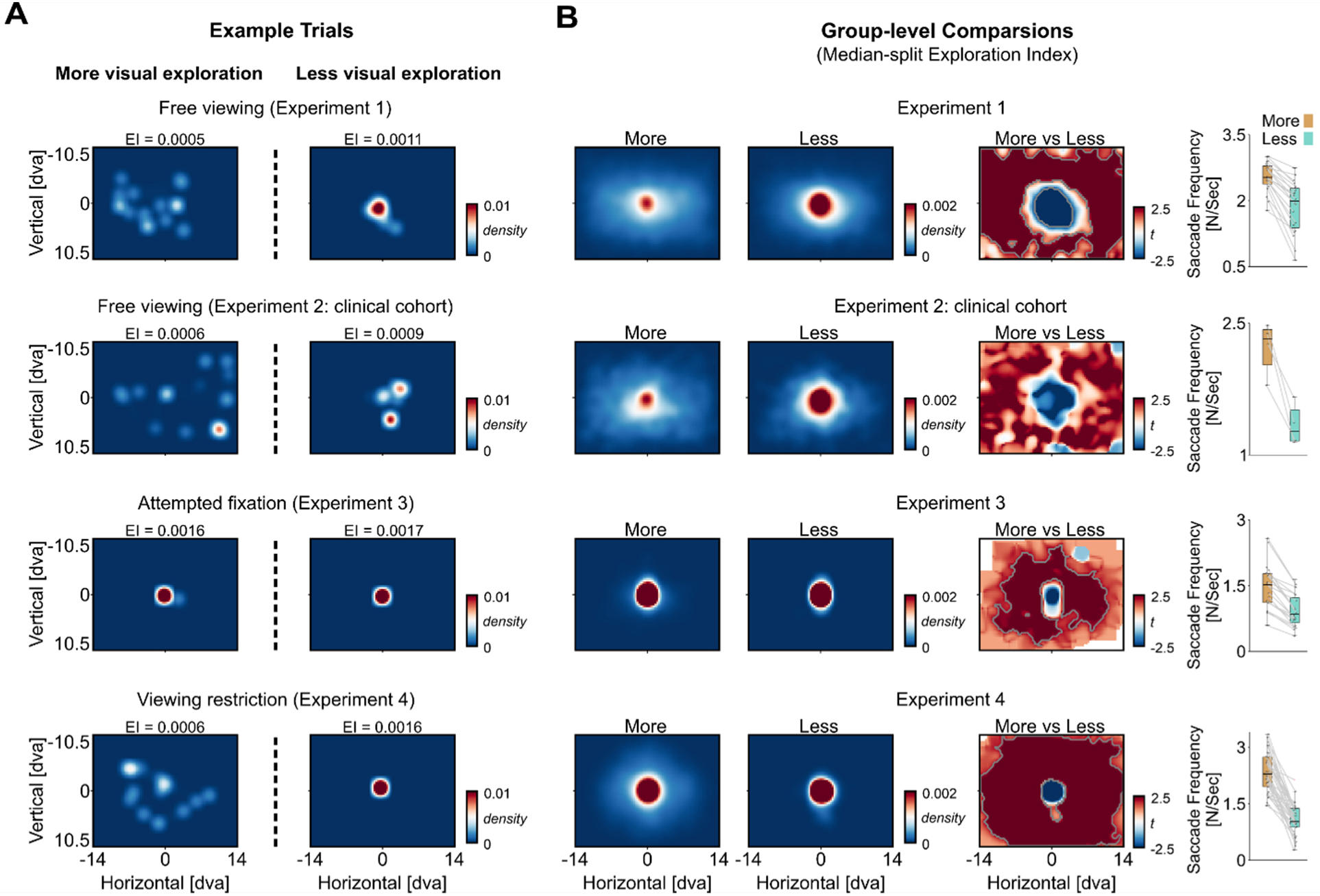
Exploration Index (EI) quantifies the degree of visual exploration. (**A**) Example trials showing 2-D gaze density arrays from Experiment 1, 2, 3 and 4 (from top to bottom). The EI is calculated as the standard deviation of the 2-D normalized gaze density array. As standard deviation measures the spread around the mean, a lower EI indicates a more uniform spread of gaze, suggesting more exploratory behaviour (left), while a higher EI reflects a peaked distribution, indicating prolonged fixation on a localized region (right). (**B**) Group-level comparisons across experiments after median-splitting trials by EI. More visual exploration (lower EI) was consistently associated with greater gaze allocations towards the surrounding locations of the scenes and less towards the centre (*p* < 0.05, two-tailed, corrected for multiple comparisons across space; note: statistical test was not applied for the clinical cohort in Experiment 2 due to low sample size, N = 4). Saccade frequency (right) followed the same patterns: trials with greater visual exploration showed higher average saccade rate across scene viewing. In all boxplots, the central mark is the median across participants and the edges of the box are the 25^th^ and 75^th^ percentiles. Grey dots represent individual data points.

**Supplementary Figure 5.**
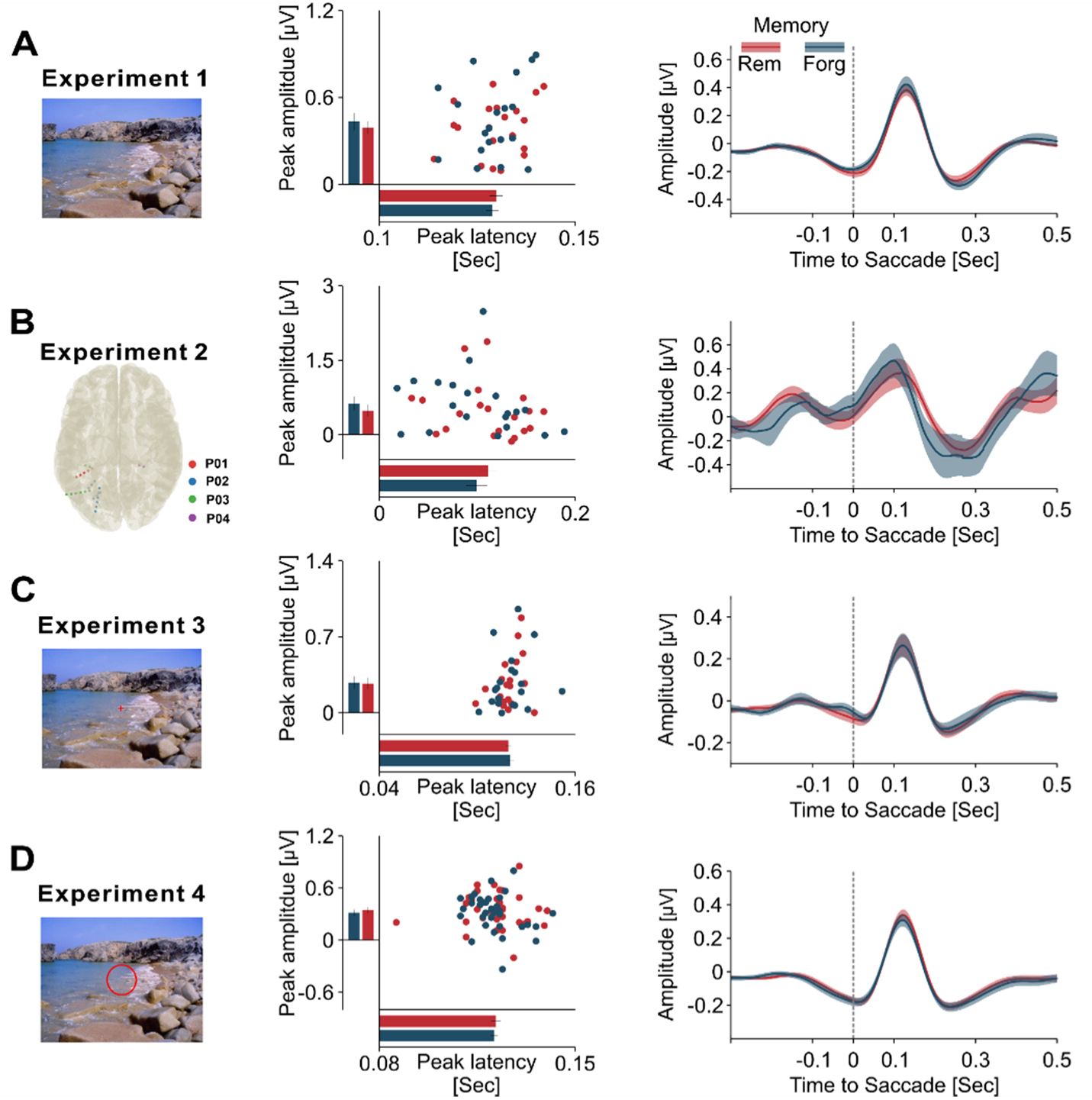
Saccade-locked alpha/beta amplitude and latency are not associated with subsequent memory performance. (**A**) During free picture viewing in Experiment 1, no significant differences in post-saccadic peak alpha/beta amplitude and latency were found between the scenes that were later remembered and forgotten. Group-level results (right) showed similar alpha/beta amplitude dynamics following saccade onset between later remembered and forgotten scenes. (**B–D**) Similar patterns were observed across different datasets: (**B**) Intracranial posterior electrodes during free viewing in Experiment 2, (**C**) Posterior scalp electrodes during attempted fixation (Experiment 3), and (**D**) Posterior scalp electrodes across varying degrees of viewing restriction (Experiment 4). Note that in (**D**), we observed a significantly higher post-saccadic peak alpha/beta amplitude for later remembered as compared to forgotten trials (remembered: M = 0.347 mV, SD = 0.202; forgotten: M = 0.316 mV, SD = 0.218, *t*_(33)_ = 2.17, *p* = 0.037), although no significant difference was found in peak latency (remembered: M = 0.122 sec, SD = 0.010; forgotten: M = 0.121 sec, SD = 0.008, *t*_(33)_ = 0.51, *p* = 0.615). Note that this effect was driven by the fact that remembered trials tended to originate from conditions with fewer viewing restriction (on average, 52.22% (SD = 12.25%) of saccades from remembered trials occurred under 21 dva condition, significantly higher than the ratio for forgotten trials: M = 39.72%, SD = 8.52%; *t*_(33)_ = 5.71, *p* < 0.001), and thus contained larger saccades (saccade size in remembered trials: M = 2.61 dva, SD = 0.67; forgotten trials: M = 2.15 dva, SD = 0.60; *t*_(33)_ = 6.27, *p* < 0.001).

**Supplementary Table 1.**
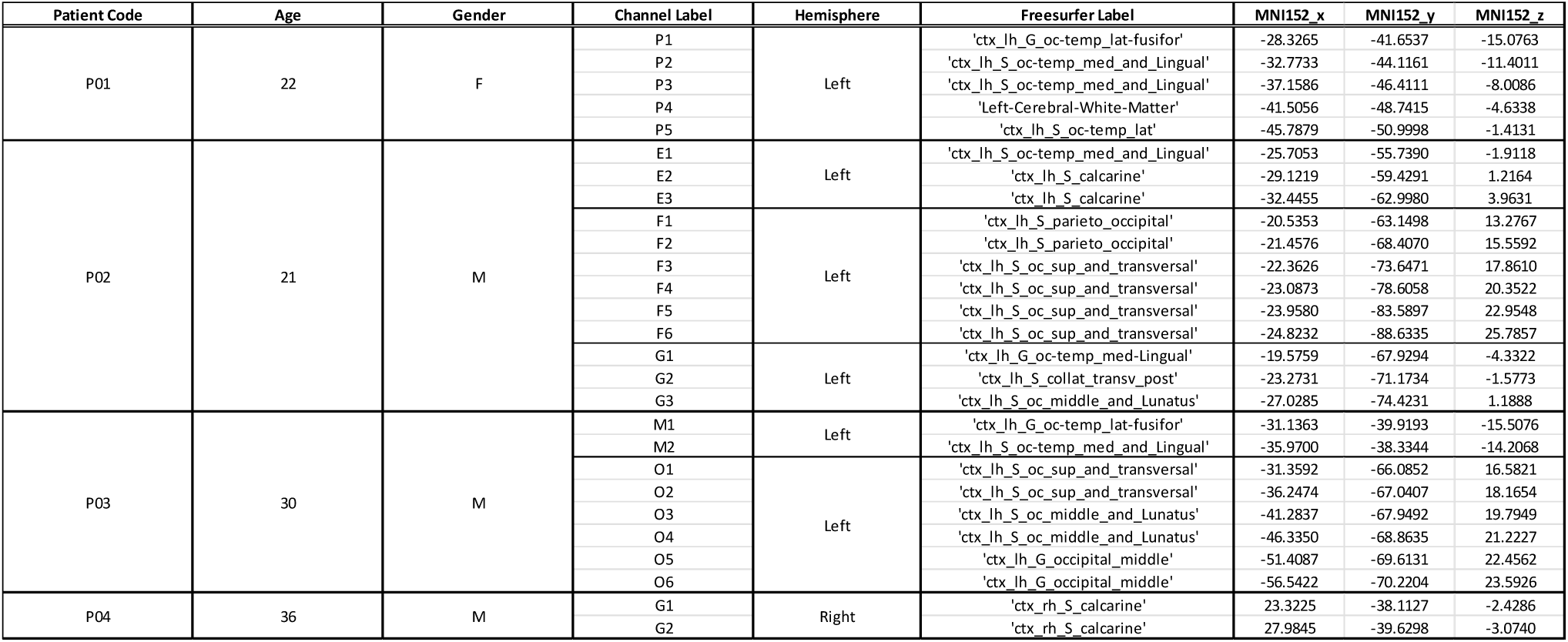
Intracranial electrode localisation. Four drug-resistant epilepsy patients with posterior electrodes (1 female; age range: 21–36 years; M = 27.25) volunteered for Experiment 2. We include the intracranial electrodes implanted in posterior brain regions into the analyses. For each patient, the electrodes’ locations in the brain were assessed with the examination of a CT and preoperative MRI T1 scans. Cerebral atlases of each patient were obtained by parcellating the preoperative T1 image using Freesurfer. The CT scan was co-registered to the corresponding T1 image, and electrode positions and anatomical labels were automatically extracted in native patient space and then manually refined using the YAEL toolbox. The table lists the hemisphere in which each electrode was placed, along with the corresponding anatomical labels and MNI (Montreal Neurological Institute) coordinates. From the 27 selected electrodes, a total of N = 20 bipolar-referenced channels were created using neighbouring contact pairs.

## Notes

### Competing Interest Statement

The authors have declared no competing interest.

### Summary of Updates

We changed the title to: Alpha/Beta oscillations reflect the dynamics of the oculomotor system: a new perspective on subsequent memory effects. We revised the time frequency analysis pipeline and updated the corresponding results figures. We additionally included FOOOF based analyses of the time frequency representations, as well as analyses of stimulus properties (measured as memorability scores) in relation to memory performance and saccade counts. These results are now reported in the supplementary materials. We also added a dedicated Methods section describing these supplementary analyses. Finally, we made several changes to the narrative in the abstract and main text, and updated parts of the discussion.

